# A commensally regulated immune rheostat fine-tunes skin barrier fitness

**DOI:** 10.64898/2026.01.07.698149

**Authors:** Anita Gola, Rohith Srinivas, Elias G. Rodig, Marina Schernthanner, Merve Deniz Abdusselamoglu, Matthew T. Tierney, Natalie J. Alexander, Kevin A. U. Gonzales, Sairaj M. Sajjath, Luis F. Soto-Ugaldi, Alain R. Bonny, Elaine Fuchs

**Affiliations:** Robin Chemers Neustein Laboratory of Mammalian Cell Biology and Development, Howard Hughes Medical Institute, The Rockefeller University; New York, USA 10065

## Abstract

At the skin’s surface, the epidermis must balance stem cell renewal with barrier maintenance to withstand environmental stress and shield against pathogens. Here, we identify a microbial-immune–epithelial feedback mechanism that integrates environmental information into stem cell regulation. Specifically, we show that Langerhans cells—an intra-epithelial macrophage population— orchestrate this circuit by producing prostaglandin E₂, which restrains stem cell proliferation, promotes epidermal differentiation and maintains barrier integrity during homeostasis. Upon pathway disruption, stem cells become overactivated, impairing differentiation and compromising barrier function. Upstream, Langerhans cell activity is tuned by the local microbial environment in a rheostat-like fashion, coupling commensal sensing to stem cell control. Our findings provide a general framework for how barrier tissues achieve adaptive homeostasis amid continual external challenge.

## MAIN TEXT

At the interface between the body and the outside world, the epidermis provides a critical, protective barrier against daily physical assaults, harmful ultraviolet radiation and skin microbes - a complex ecosystem composed of bacteria, archaea, fungi and viruses (*1, 2*). To withstand these insults, the mammalian epidermis is a stratified epithelial tissue that undergoes continual turnover. The drivers for these dynamics are the epidermal stem cells (EpSCs) that reside in the basal layer, nestled beneath the skin’s outermost stratified layers and anchored atop the basement membrane of specialized extracellular matrix that separates the epidermis from dermis (Fig. 1A) (*3*). Functionally, EpSCs proliferate and differentiate, generating a tightly regulated upward flux of suprabasal progeny that terminally differentiate to form a protective barrier of dead, flattened squames that are then shed from the skin surface and replaced by differentiating cells moving outward.

**Figure. 1:**
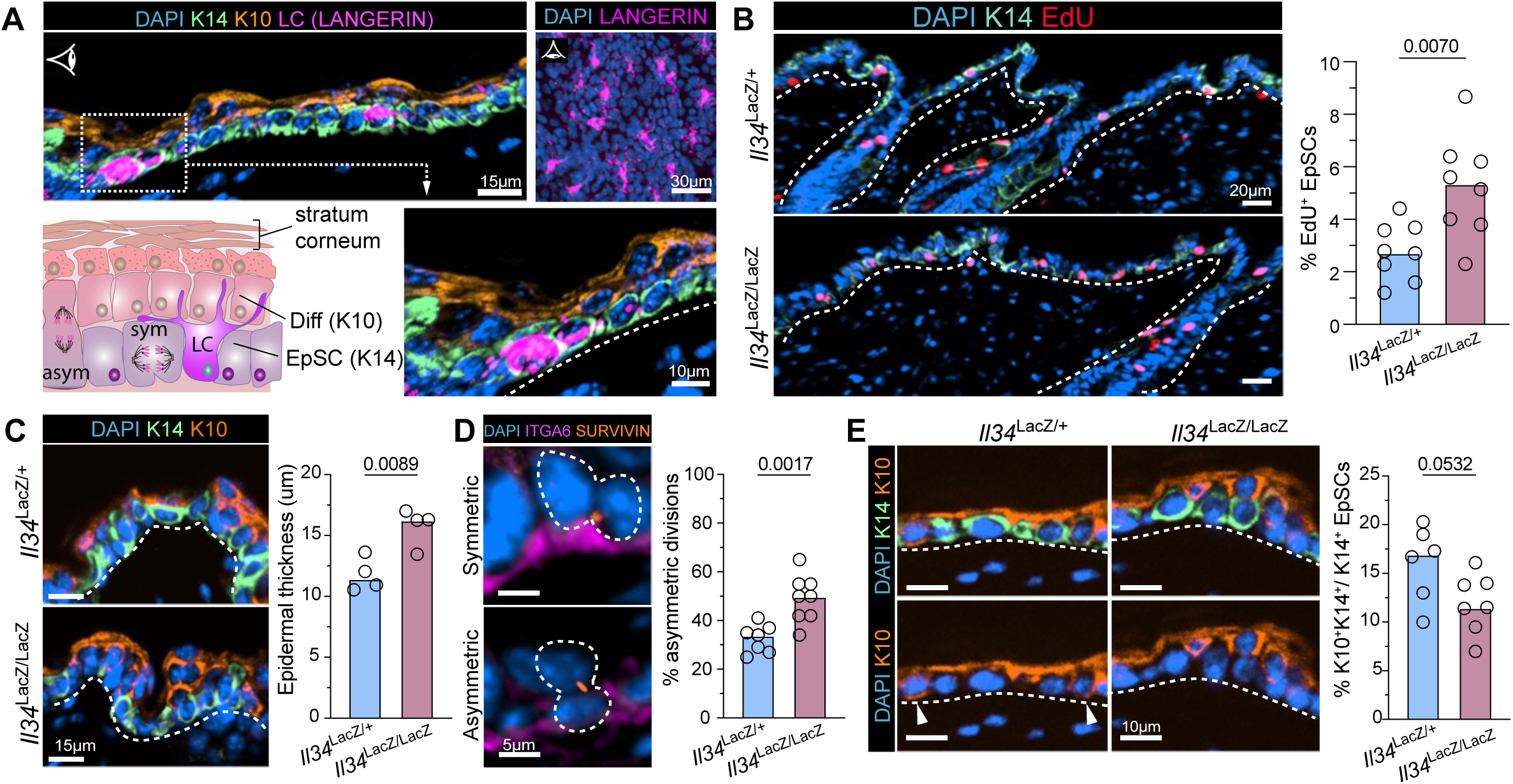
Loss of LCs results in altered EpSC function. **(A)** Representative schematic and immunofluorescent (IF) image of postnatal day P60 epidermis showing the EpSC compartment in the basal layer (marked by K14), committed differentiating progeny (K10) and Langerin-expressing LCs. Boxed area is magnified in lower right. Dashed line denotes epidermal-dermal border. Schematic eyes denote sagittal vs planar orientations of sections. **(B)** Representative IF images and quantifications of EpSC proliferation (S-phase EdU labeling) in control (*IL34^LacZ/+^*) vs LC-deficient (*IL34^LacZ/LacZ^*) skin. **(C)** Representative IF images and quantifications showing increased epidermal thickness upon LC loss. **(D)** Representative IF images of division planes of anaphase EpSCs as marked by Survivin and DAPI, relative to the basement membrane, proxied by ITGA6. Quantifications of asymmetric divisions (defined as one daughter being nonadjacent to the basement membrane) relative to total divisions within the basal layer of EpSCs. **(E)** Representative IF images and quantifications of the percentage of committed K10^+^ K14^+^ double-positive out of the total K14^+^ pool of basal layer progenitors. For (A-E): Each dot corresponds to data from one mouse with median shown, n ≥ 4 mice per condition, two-tailed unpaired Student’s t tests.

Although the major pathways and transcriptional programs governing epidermal differentiation have been well characterized, the inhibitory processes that dampen EpSC activity to preserve homeostasis are still poorly understood (*3–5*). Additionally, while EpSCs are protected from direct environmental exposure by the stratified layers above, this insulation limits their ability to directly sense and respond to changes at the skin surface. How EpSCs receive and integrate environmental information has remained elusive and yet is critical for maintaining an optimal homeostatic equilibrium. The problem is particularly daunting considering that at any given moment, the basal layer of EpSCs must simultaneously coordinate the cell divisions that preserve a functional pool of stem cells with those that produce committed progeny tasked with rejuvenating the body surface.

One way for biological systems to maintain robustness and adaptability while facing external perturbations is in the implementation of negative feedback control (*6*). In the context of stem cell biology, these regulatory signals often derive from the neighboring “niche”– the cellular players and physical micro-environment that regulate stem cell fate. In the murine epidermis, there are only two populations of neighboring resident cells that co-exist with EpSCs and might serve in this capacity: Dendritic Epithelial T Cells (DETCs), a type of γδ T lymphocyte; and Langerhans cells (LCs), a specialized tissue-resident macrophage. Unlike DETCs, LCs are uniquely found in all types of stratified epithelia with high barrier demands (*7, 8*), and are conserved across the mammalian kingdom (*9, 10*). These are important features if they are to play a generalizable role in regulating stem cell function in these tissues. In addition, LCs are highly sensitive environmental sensors, able to constantly sample their surroundings and extend their dendrites through the stratified layers of the barrier epithelium (*8, 10, 11*).

The ability of LCs to take up antigens from their environment and initiate adaptive immune responses has resulted in their characterization as potent antigen-presenting cells (*10, 12, 13*). Yet, they reside in constant contact with EpSCs for large periods of time, placing them in an ideal position to potentially influence stem cell behavior. Despite this proximity, their role in maintaining epidermal homeostasis and regulating EpSC output has remained largely unexplored. With the additional consideration that other types of tissue-resident macrophages have been implicated in stem cell regulation in various tissues and contexts (*14–17*), we turned our attention to the possibility that LCs might be able to relay critical environmental information to fine-tune EpSC function.

Here, we tackle whether and how this highly abundant resident immune cell population functions within the epidermis during homeostasis. By examining their local function within the EpSC niche, we unearth a hitherto unrecognized role for LCs in modulating EpSC dynamics and promoting barrier fitness. Specifically, we identify a novel circuit in which LCs regulate EpSC proliferation and promote skin barrier fitness through the production of prostaglandin E₂ (PGE₂). Further, we find that the local microbiota regulates this crosstalk by tuning in a rheostat-like fashion PGE₂ levels, and consequently EpSC function. This circuit aligns EpSC output with the demands of its microbiota-shaped microenvironment, shedding new light on the mechanisms that govern tissue homeostasis.

### LC deficiency perturbs EpSC function

To decipher the role of LCs in epidermal biology, we began by leveraging the previously generated IL-34 deficient animals (*Il34*^LacZ/LacZ^) known to have radically reduced LCs (fig. S1A) (*18, 19*). In contrast to models like huLang-DTR mice (*10*), these animals lack LCs from birth, and cannot be compensated by other monocyte-derived macrophages, because the epidermis lacks the critical trophic factor that supports macrophage growth and maintenance (fig. S1A-B). This distinction is important because acute, synchronized LC ablation in adult mice would abruptly create physical space through LC death, which could confound interpretation by triggering EpSC proliferation and low-grade inflammation.

Despite avoiding such caveats, we observed that ∼ 50% more EpSCs were in S-phase in *Il34* ^LacZ/LacZ^ animals compared to *Il34* ^LacZ/+^ littermates, as judged by administering a 4-hour pulse of the nucleotide analogue 5-Ethynyl-2’-deoxyuridine (EdU) to adult (P60) animals prior to analyses (Fig. 1B). This surprising result was accompanied by a small but significant increase in epidermal thickness as quantified by immunofluorescence using the EpSC marker keratin K14 and the early differentiation marker keratin K10, and as observed in histological Hematoxylin and Eosin (H&E) sagittal skin sections (Fig. 1C and fig. S1C). Additional histological features included a subtle change in epidermal morphology, with basal EpSCs more columnar in shape when compared to littermate controls. Consistent with these features was an increase in the number of EpSC divisions that occurred perpendicularly to the underlying basement membrane (Fig. 1D). Such divisions have been observed concomitant with stratification in early skin development and have been linked as a means of balancing basal epidermal density (*20, 21*). Lastly, we investigated whether EpSC differentiation was altered in these animals and found that the frequency of early committed, K10/K14^+^ cells in the basal layer was decreased in animals that lacked LCs (Fig. 1E). While modest, these differences pointed to a view that the loss of LCs shifts the normal balance between EpSC quiescence, proliferation and differentiation that exists in homeostasis, alluding to a loss of feedback control.

### A Prostaglandin Signaling Axis connects LCs and EpSCs in the Skin

To dissect the underlining mechanism by which LCs influence EpSCs during skin homeostasis, we assembled a comprehensive single-cell RNAseq dataset from in-house and publicly available data enriched in immune and epithelial cells (see *Methods*) (*22*). Unsupervised clustering of the combined samples identified 13 clusters, which we visualized using Uniform Manifold Approximation and Projection (UMAP) (Fig. 2A). We first asked which genes distinguish LCs from the other clusters. We were struck by *Ptgs1,* encoding the enzyme cyclooxygenase 1 (COX-1), which among the resident immune cell populations in the skin was exclusively expressed by LCs (Fig. 2B, fig. S2A). Transcripts were also detected in EpSCs and differentiated (Diff) cells, but at markedly lower levels than LCs, a finding that was also observed by intra-cellular COX-1 staining in flow cytometry (Fig. 2C, fig. S2B-C). Further, by immunofluorescence microscopy, COX-1 displayed a striking co-compartmentalization with Langerin, the canonical LC marker (Fig. 2D).

**Figure. 2:**
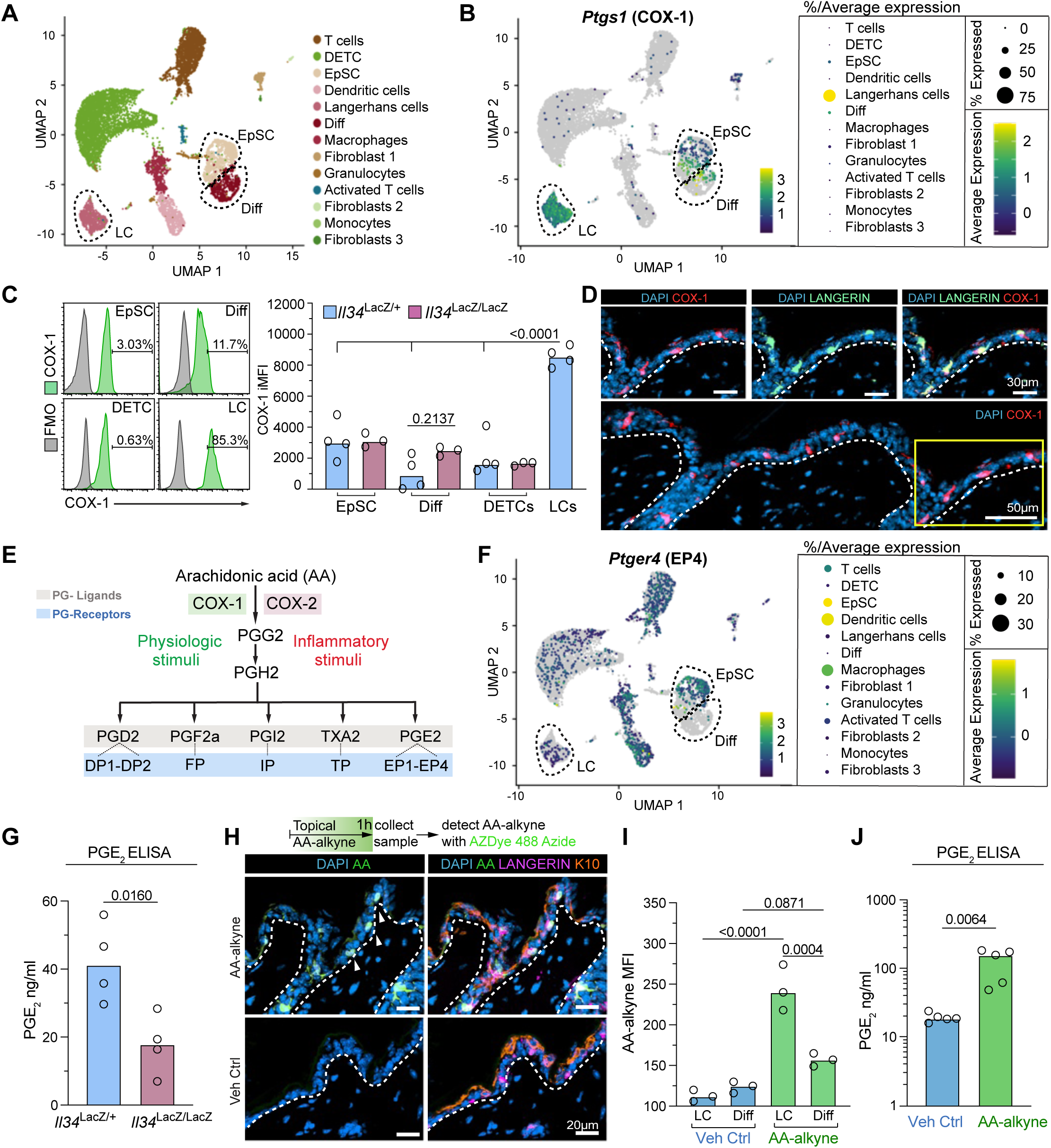
COX-1 expression is uniquely enriched in epidermal Langerhans cells. **(A)** UMAP plots of murine skin single cell RNA-seq data encompassing 9736 cells (7897 immune cells (thereof 695 Langerhans cells), 1535 epithelial cells (composed of 890 EpSCs and 645 differentiated progeny), 304 fibroblasts, respectively). Each cell is colored according to cell type annotation as described in legend (see Methods). **(B)** UMAP plot (left), average relative expression and percentage of *Ptgs1^+^* cells within the skin populations. **(C)** Left: Representative histogram from flow cytometry of COX-1 expression by EpSCs, differentiating epidermal cells (Diff), DETCs and LCs within the epidermis of control (*Il34*^LacZ/+^) animals. Right: Integrated Mean Fluorescence Intensity (iMFI) of COX-1 in EpSCs, Diff, DETCs and LCs from control and LC-deficient animals (*Il34*^LacZ/+^ and *Il34*^LacZ/LacZ^ respectively). Each dot corresponds to data from one mouse, median shown, One-Way ANOVA with Šidák multiple-comparison correction. **(D)** Representative IF image and zoom-in insets of COX-1 expression within the skin, showing enrichment of COX-1 signal within LCs. **(E)** Prostaglandin pathway schematic highlighting COX-1 and COX-2 enzymes in the production of prostaglandins (PG) (highlighted in grey), with cognate receptors (PG-R) (highlighted in blue). **(F)** UMAP plot (left), average expression and frequency of total of *Ptger4+* cells within the skin populations. **(G)** PGE_2_ levels within epidermal fractions of control and LC-deficient animals (*Il34*^LacZ/+^ and *Il34*^LacZ/LacZ^ respectively), two-tailed unpaired Student’s t tests. **(H-I)** Top: Schematic of arachidonic acid (AA)-alkyne topical application following detection with AZDye-488 Azide via click-chemistry, Bottom: representative IF image of AA uptake by LCs, with respective quantification of AA-alkyne Mean Fluorescent Intensity (MFI), One-Way ANOVA with Šidák multiple-comparison correction. **(J)** PGE_2_ levels within epidermal fractions of WT animals in the presence or absence of AA-alkyne, two-tailed unpaired Student’s t tests. For (A-J): Each dot corresponds to data from one mouse with median shown, n ≥ 3 mice per condition.

COX enzymes are central in the biosynthetic pathway for hormone-like lipid mediators called prostaglandins, which exert highly localized effects due to their rapid degradation (Fig. 2E) (*23*). Induced by inflammatory cues, COX-2 has been extensively studied in the context of wounding and inflammation, where its upregulation drives the production of prostaglandins that activate immune cells, increase blood flow, and sensitize pain neurons (*23*). By contrast, COX-1 is generally considered to be constitutively expressed and to support the maintenance of normal physiological function. However, COX-1’s functional relevance in regulating the epidermis during skin homeostasis remains largely unexplored (*24–31*), and the molecular mechanisms by which COX-1–derived prostaglandin signaling influences EpSC function are poorly understood. Notably, the striking enrichment of COX-1 expression in LCs suggests that its localization is unlikely to be incidental and instead points to a specialized, context-dependent role beyond general housekeeping.

Based upon the evidence at hand, we posited that prostaglandins might be functioning in LC:EpSC crosstalk in a manner that might be integral to controlling skin homeostasis. If prostaglandins generated by LCs directly signal to affect skin epithelial cells, they should express one or more of the known prostaglandin receptors. To this end, the only prostaglandin receptor we detected in EpSCs was *Ptger4,* encoding the G-protein coupled receptor Prostaglandin E_2_ receptor 4 (EP4), which is specific for prostaglandin E_2_ (PGE_2_) (Fig. 2F, fig. S3A-D). In line with the spatial vicinity of EpSCs and LCs to the epidermal basal layer, Cell-chat ligand–receptor analyses further supported prostaglandin signaling as a likely axis through which these two cell populations communicate (fig. S3E).

To draw a physiological link between COX-1 expression by the LCs and prostaglandin signaling in the EpSCs, we assessed if LCs can directly make PGE_2_. We first showed by ELISA analysis that loss of LCs within the epidermis was associated with decreased PGE₂ abundance (Fig. 2G). Conversely, we hypothesized that if LCs represent the principal source of PGE₂ in the homeostatic epidermis, then substrate supplementation to LCs should elicit measurable PGE₂ synthesis. In devising a method to test this possibility, we reasoned that by topically applying an alkyne-tagged version of the prostaglandin precursor arachidonic acid (AA), we might be able to use click chemistry to track the uptake of COX-1’s substrate. If so, this would provide us a means to directly assess PGE₂ production by LCs. Indeed, following topical application, click chemistry of AA-alkyne uptake showed internalization by LCs in the epidermis (Fig. 2H-I), which was accompanied by a marked increase in PGE₂ levels (Fig. 2J). Thus, not only are LCs the major producer of COX-1 within the epidermis, but they also harbor the capacity to be the principal generator of PGE₂ within the niche.

### LC-derived PGE2 plays a critical role in directly regulating epidermal homeostasis

To understand the specific effects of PGE_2_ sensing on epidermal stem-cell function, we first took a pharmacological approach to modulate prostaglandin levels within homeostatic skin. We inhibited the prostaglandin pathway by topically applying indomethacin (Indo), a COX nonselective cyclooxygenase inhibitor; iCOX-1, a COX-1–selective inhibitor; and iEP4, an EP4 receptor antagonist. Conversely, we raised endogenous levels of PGE_2_ by topically applying a) a stable derivative of PGE_2,_ 16,16-dimethyl PGE_2_ (dmPGE_2_), and b) an inhibitor of the key degrading enzyme for PGE₂, 15-PGDH (15-hydroxyprostaglandin dehydrogenase) (*32*). In all permutations, EpSC proliferation was altered—increasing with prostaglandin inhibition and decreasing with prostaglandin augmentation (Fig. 3A-B, fig. S4A).

**Figure 3.**
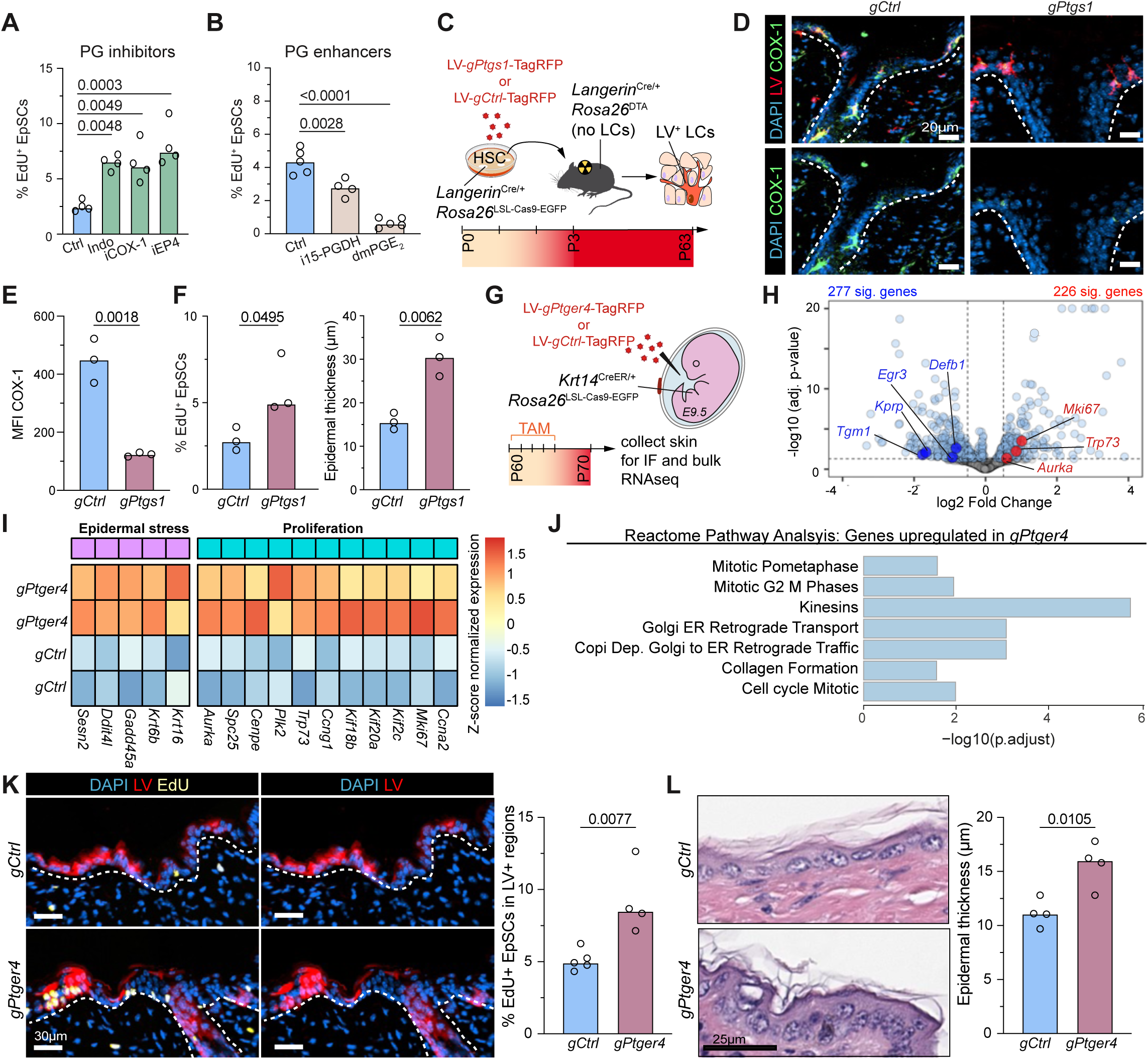
Loss of EP4 signaling in EpSCs increases stem cell activity. (A-B) EpSC proliferation (S-phase EdU labeling) following 24h pharmacological application of PG inhibitors [Indo (Indomethacin; COX non-selective inhibitor), iCOX-1 (SC-560; selective COX-1 inhibitor), iEP4 (ONO-AE3-208; EP4 receptor antagonist)] or PG enhancers [i15-PGDH (SW033291; 15-PGDH inhibitor), dmPGE_2_]. One-Way ANOVA with Šidák multiple-comparison correction. **(C)** Experimental schematic for genetic perturbation of LCs. **(D-E)** Representative IF image and COX-1 MFI of LCs from LV expressing *gCtrl* or *gPtgs1.* **(F)** EpSC proliferation (S-phase EdU labeling) and epidermal thickness in *gCtrl* or *gPtgs1* animals. **(G)** Experimental schematic of *in utero* LV delivery for genetic perturbation of EpSCs. **(H)** Volcano plot of differential gene expression between *gCtrl* or *gPtger4* EpSCs. Genes associated with proliferation, differentiation and barrier function highlighted. **(I)** Heatmap showing relative Z-score normalized expression of key genes related to epidermal stress and proliferation in *gPtger4* or *gCtrl* EpSCs. **(J)** Pathway-level analysis using Reactome database in upregulated genes in *gPtger4* EpSCs. **(K)** Representative IF images and quantification of EpSC proliferation (S-phase EdU labeling) in *gCtrl* or *gPtger4* LV^+^ regions. **(L)** Representative H&E staining and quantifications of epidermal thickness in *gCtrl* or *gPtger4* (transduced with 70% efficiency). For (A-L): Each dot corresponds to data from one mouse with median shown, n ≥ 3 mice per condition. Unless otherwise stated: two-tailed unpaired Student’s t tests.

To investigate the importance of LC-derived PGE₂ we developed a novel strategy to ablate *Ptgs1* specifically in LCs (Fig. 3C). Leveraging the fact that LCs are repopulated by monocyte-bone-marrow derived macrophages from hematopoietic stem cells (HSCs) (*33, 34*), we cultured HSCs from *huLangerin*^Cre/+^; *Rosa26*^LSL-Cas9-EGFP^ animals in the presence of lentiviruses harboring either Cas9 *Ptgs1* or *Ctrl* targeting sgRNAs (*see* Methods). Following selection for LV^+^ HSCs based upon TagRFP expression, we transferred the cells into γ-irradiated *huLangerin*^Cre/+^; *Rosa26*^DTA^ animals which lack LCs in the epidermis (Fig. 3C). Two months after adoptive transfer, efficient LC reconstitution had occurred within the skin epidermis of *gPtgs1* and *gCtrl* animals (at approx. 70% reconstitution when compared to WT animals) (Fig. 3D, fig. S4B). Affirming our strategy, COX-1 expression was selectively reduced in LCs transduced for *gPtgs1* and not *gCtrl* (Fig. 3D-E). In accordance with our pharmacological modifications, animals with COX-1 deficient LCs exhibited increased EpSC proliferation and epidermal thickening (Fig. 3F, fig. S4C).

To test whether LC-derived PGE_2_ directly affects EpSC prostaglandin receptor signaling, we turned to a cell type-specific genetic approach. We leveraged *in utero* lentiviral delivery of *Ptger4* targeting sgRNAs, into the amniotic sacs of *Krt14*^CreER/+^; *Rosa26*^LSL-Cas9-EGFP^ E9.5 mouse embryos (Fig. 3G) (*35*). At this stage, the single layer of embryonic skin epithelial progenitors becomes selectively and stably transduced (*35*). Following tamoxifen application in adulthood, Cre recombinase then becomes activated primarily in the EpSCs, enabling us to target ablation of *Ptger4* specifically in the epidermal keratinocytes (fig. S4D), and confirm that the interruption in PGE_2_ signaling within EpSCs did not affect LC numbers (fig. S4E).

To investigate how loss of EP4 signaling affected the epidermis, we purified EpSCs by fluorescence activated cell sorting (FACS) for (CAS9) eGFP, antibodies against surface integrins α6 and β1, and (LV^+^) TagRFP, and subjected them to RNA sequencing. Differential gene expression analysis of EP4-deficient EpSCs revealed broad transcriptional shifts, with hundreds of genes significantly upregulated ≥2X (p<0.05) compared to controls (Fig. 3H). Using Reactome, a manually curated database of biological pathways and processes that are organized as a network of reactions, we observed a strong enrichment for processes linked to stress responses and epithelial proliferation (Fig. 3I-J). These signatures were bolstered by an increase in EdU incorporation within the basal layer of *Ptger4-*null epidermis, accompanied by an increase in epidermal thickness (Fig. 3K-L). In addition, immunofluorescence microscopy revealed activation of the stress response keratin, K6 within the *Ptger4-*null epidermis (fig. S4F). Together, these results suggested that loss of prostaglandin signaling in EpSCs perturbs the brake on stem cell self-renewal.

### Epithelial prostaglandin EP4 receptor signaling sustains EpSC quiescence by inhibiting YAP nuclear translocation

To gain deeper insights into how EP4 signaling impacts tissue homeostasis, we considered three previously described regulatory pathways, namely CREB, β-catenin/WNT and YAP signaling (*36–39*), which have been described in other contexts to be altered when EP4 signaling is perturbed (Fig. 4A). However, upon revisiting our bulk RNA-seq dataset and performing Gene Set Enrichment Analysis (GSEA) to calculate enrichment scores, we only saw evidence of altered target gene expression for YAP signaling upon loss of EP4 and not for β-catenin/LEF/TCF or CREB (Fig. 4B). Consistently, immunofluorescence microscopy revealed a marked increase in nuclear YAP within the *Ptger4* null epidermal patches where EP4 was absent (Fig. 4C), a phenotype that was similarly observed in animals lacking LCs (fig. S4G). This was particularly intriguing given the well-established role for nuclear YAP/TEAD signaling in promoting proliferation within the basal epidermal layer (*40, 41*).

**Figure 4.**
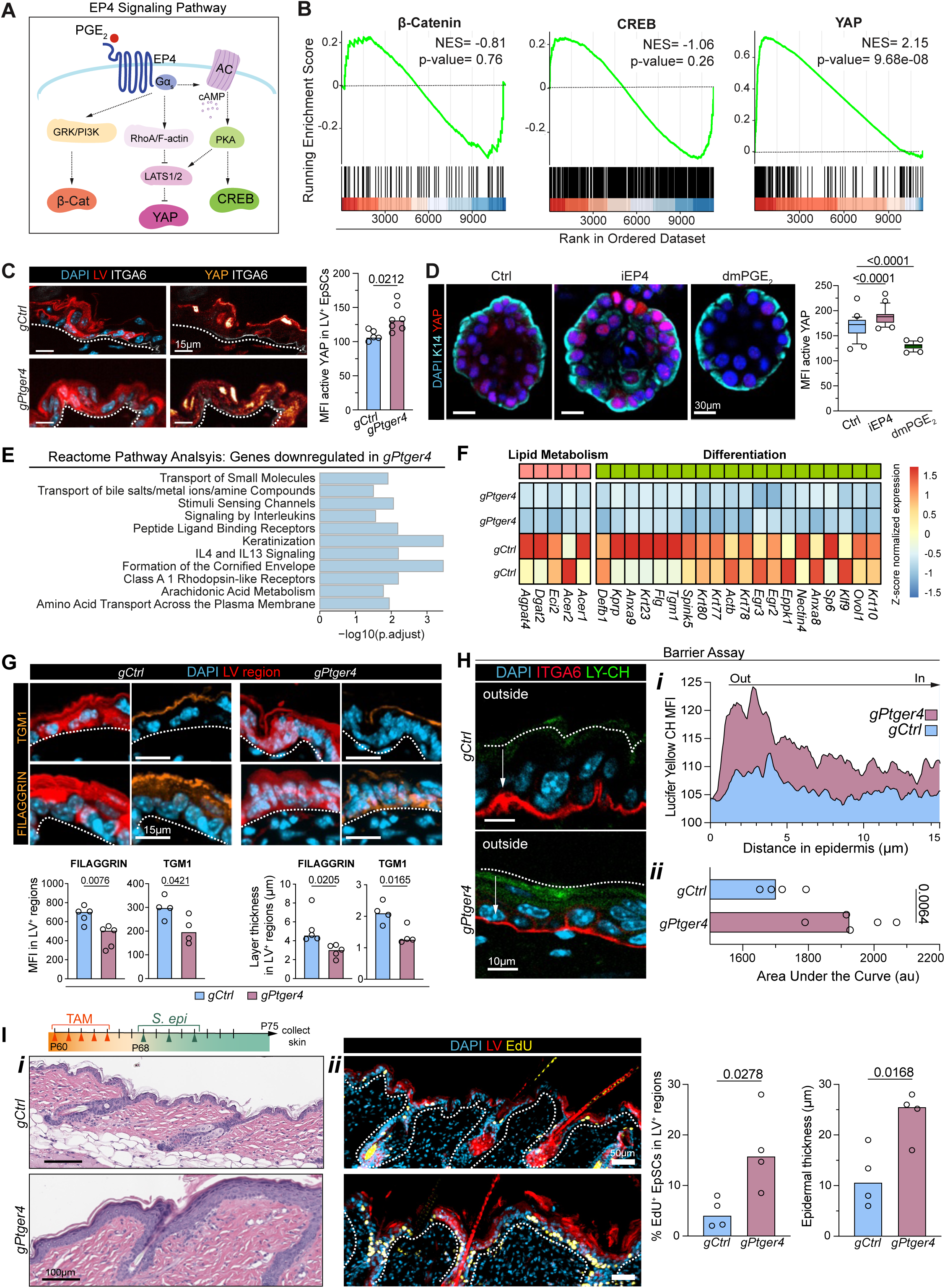
EP4-deficient EpSCs exhibit increased YAP activation and compromised barrier integrity with heightened responses to commensal stimulation. **(A)** Schematic of EP4 signaling pathway. **(B)** GSEA running enrichment score for β-catenin, CREB, and YAP pathways, showing that only YAP correlates with loss of EP4 signaling in EpSCs. **(C)** Representative IF images and MFI quantifications of nuclear active YAP (shown as an intensity heatmap) in EpSCs from *gPtger4* or *gCtrl* LV^+^ regions as demarked by ITGA6 expression. **(D)** Representative IF images and MFI quantifications of nuclear active YAP in Ctrl, iEP4, and dmPGE_2_ treated organoids. Each dot represents an organoid; Box-plot: 10-90 percentile, One-Way ANOVA with Šidák multiple-comparison correction. **(E)** Pathway-level analysis using Reactome database in down regulated genes in *gPtger4* EpSCs. **(F)** Heatmap showing relative Z-score normalized expression of key genes related to lipid metabolism and differentiation in *gPtger4* or *gCtrl* EpSCs. **(G)** Top: Representative IF images of TGM1 and FILAGGRIN staining in *gCtrl* or *gPtger4* treated animals. Bottom: MFI quantifications of stated markers, and layer thickness in LV^+^ region of *gCtrl* or *gPtger4* sagittal sections. **(H)** Barrier Assay: Representative IF images of Lucifer Yellow CH (Ly-CH) following topical application in *gCtrl* or *gPtger4* animals (dotted line denotes outside border of epidermis); i) Representative Ly-CH MFI trace from *gCtrl* or *gPtger4* animals as measured from the outside of the epidermis to the epidermal-dermal boundary as demarked by ITGA6 expression; ii) Area under the curve of Ly-CH MFI expression in *gCtrl* or *gPtger4* animals. **(I)** Experimental schematic, representative H&E stained sagittal sections (i), and IF images of EdU incorporation (ii) and respective quantifications of EpSC proliferation (S-phase EdU labeling) and epidermal thickness in *gCtrl* or *gPtger4* animals following *S.epi* challenge. For (A-I), unless otherwise stated: dotted lines denote epidermal-dermal border; each dot corresponds to data from one mouse; n ≥ 3 mice per condition; median shown; two-tailed unpaired Student’s t tests.

To assess whether the effects of *Ptger4* loss of function were directly linked to activation of YAP, we turned to EpSC organoid cultures, where we could manipulate the EpSC prostaglandin levels independently of LCs or other cell populations (fig. S5A) (*42*). After using qPCR to confirm expression of *Ptger4* and *Ptgs1* in organoids (fig. S5B), we pharmacologically inhibited EP4 signaling to inhibit organoid-intrinsic PGE_2_ sensing and conversely increased EpSC prostaglandin signaling levels by adding dmPGE_2_ to the culture medium. These pharmacological manipulations recapitulated the *in vivo* phenotype seen upon LC immune cell loss, yielding enhanced EpSC proliferation, larger organoid colonies and nuclear YAP activity when PGE_2_ signaling was restricted and a suppression of these features when PGE_2_ was stabilized (Fig. 4D, fig. S5C). Further, we found that PKA activity (known to be a direct second messenger downstream of Gα-coupled receptors that can modulate LATS (*37, 43*)) exhibited a robust dose-dependent activation following *in vitro* stimulation of EpSCs with dmPGE₂ (fig. S5D). Taken together, these data suggested that EP4 activity in EpSCs functions to maintain quiescence within the stem cell compartment, at least in part by inhibiting the Hippo-YAP pathway.

### Loss of epithelial EP4 signaling results in compromised barrier fitness

Previous work showed that when mice are genetically engineered to hyperactivate YAP in the skin epidermis, elevated proliferation occurs concomitantly with repression of differentiation (*40, 41*). To more rigorously probe the possible link between loss of PGE₂ signaling and YAP activation, we looked at the downregulated genes in our *Ptger4* loss of function transcriptome. Indeed, Reactome Pathway Analysis for downregulated genes included formation of the cornified envelope, keratinization, fatty acid metabolism and immune cell signaling (Fig. 4E). Examination of these transcripts revealed a marked downregulation of *Krt10* and *Ovol1,* encoding early differentiation markers; *Flg* and *Tgm1*, encoding proteins required to assemble the skin barrier; and *Acer1* and *Eci2*, involved in forming the lipid bilayers that seal the dead keratinocyte squames at the skin surface (Fig. 4F). The changes were also reflected *in vivo* and *in vitro* by immunofluorescence for several components of late differentiation, including Filaggrin (FLG), and Transglutaminase 1 (TGM1), all showing a decrease both in mean fluorescence intensity and overall thickness when EP4 signaling was perturbed (Fig. 4G, fig. S5E). In addition, the frequency of K10/K14^+^ cells in the basal layer was subtly decreased in *Ptger4* targeted regions *in vivo*, and in organoids, all consistent with the previously reported altered differentiation program in YAP over-activation (fig. S5F-G) (*40, 41*).

To functionally investigate whether altered prostaglandin signaling during epidermal homeostasis compromises the skin’s barrier, we performed a barrier permeability assay in animals injected with LV-*gPtger4* or LV-*gCtrl*. Topical Lucifer Yellow carbohydrazide salt (LY-CH) application revealed increased epidermal penetration, consistent with an impaired outside-in barrier integrity that depends upon an intact granular and stratum corneum (Fig. 4H). Lastly, we put the epidermal barrier to the test by challenging it with *Staphylococcus epidermidis* (*S. epi*), a well-established gram-positive bacterial model of skin commensalism in humans (*47, 48*). Consistent with prior reports on wild-type mice (*48, 49*), when the skin barrier was intact, *S. epi* did not elicit overt skin pathology or inflammation. However, in *Ptger4*-null regions, the epidermis reacted strongly to *S. epi* application, with increased EpSC proliferation and epidermal hyperthickening, characteristic of a pathological skin response (Fig. 4I). A similar phenotype was also observed following *S. epi* challenge in COX-1 LC deficient animals, as well as animals that were concurrently treated daily with iCOX-1 (fig. S6A-D). Altogether, these data suggested that when the LC-EpSC circuit governing prostaglandin signaling is compromised, all-encompassing changes to skin barrier function arise.

### Skin Commensals Fine-Tune COX-1 Expression in a rheostat-like fashion

Considering the importance of PGE_2_ signaling in EpSCs for optimal barrier fitness, and having uncovered the underlining molecular mechanism, we turned our attention to understanding what regulates PGE_2_ production in the epidermis. Although often regarded as a pro-inflammatory mediator, PGE₂ plays a more nuanced role in homeostasis, with suggested anti-inflammatory properties and contribution to lipopolysaccharide (LPS) tolerance (*50–53*). As our findings pointed to COX-1-expressing LCs as a central producer of PGE_2_ in the epidermis, we focused our attention on what might be regulating *Ptsg1* in these cells. Given the well-established role of LCs as immune sentinels, we hypothesized that the skin microbiota might directly modulate COX-1 expression in these cells because of their intimate contact with the epidermal environment.

To test this hypothesis, we measured COX-1 levels in LCs when mice were raised in a germ-free (GF) environment and compared them with those housed in normal Specific Pathogen Free (SPF) conditions. Additionally, we compared COX-1 levels in LCs from homeostatic SPF skin that had been colonized with *S. epi*. As judged by mean fluorescence intensity, COX-1 protein was markedly diminished in the LANGERIN+ LCs of germ-free animals when compared to SPF controls, and conversely, it rose dramatically in *S.epi-*colonized animals (Fig. 5A). Since LCs were the major producers of COX-1, we could use RT-qPCR of mRNA purified from bulk epidermal tissue as a proxy to verify this trend at the transcriptional level (Fig. 5B).

**Figure 5.**
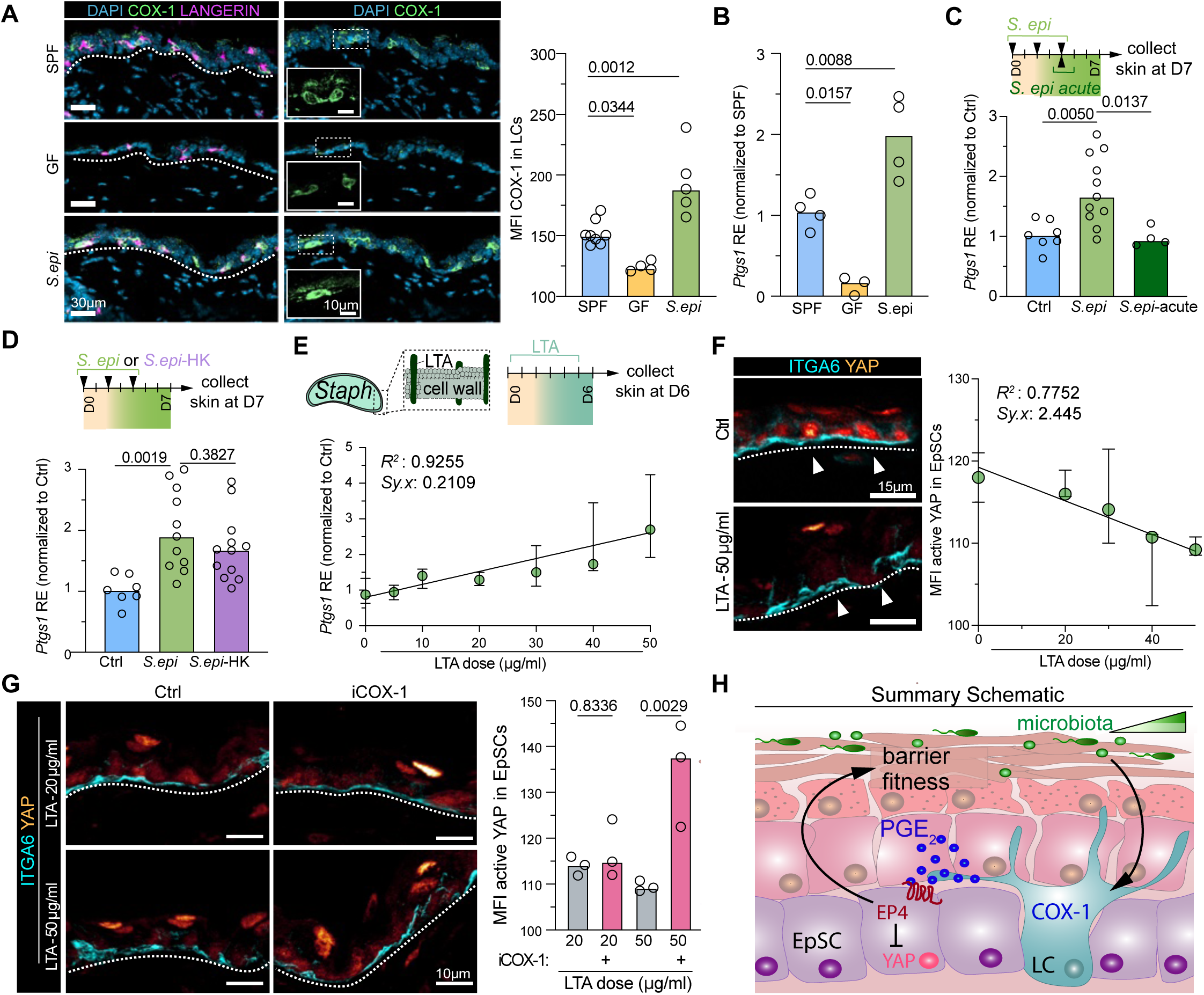
Commensal cues modulate COX-1 expression in LCs, driving proportional changes in EpSC activity. **(A)** Representative IF images and mean fluorescence intensity (MFI) quantifications of COX-1 in LCs of mice housed under SPF, GF or *S.epi-*colonized conditions. **(B)** Epidermal Relative Expression (RE) of *Ptgs1* in the skins of mice housed under SPF, GF or *S.epi-*colonized conditions (normalized to SPF animals). **(C)** Top: Experimental schematic highlighting *S.epi* application regimen; Bottom: epidermal relative expression of *Ptgs1* in Ctrl, *S.epi, S.epi-acute* colonized animals (normalized to Ctrl animals). **(D)** Top: Experimental schematic highlighting *S.epi* and *S.epi* – heat killed (*S.epi* – HK) application regimen; Bottom: epidermal relative expression of *Ptgs1* in Ctrl, *S.epi, S.epi-acute* colonized animals (normalized to Ctrl animals). **(E)** Top: Experimental schematic highlighting the topical application regimen for LTA, as well as its location within the bacterial cell wall; Bottom: epidermal relative expression of *Ptgs1* according to LTA dose. Median with Standard Error (SE) shown, simple linear regression fitted. **(F)** Representative IF images (dashed line epidermal-dermal boundary, arrowheads highlighting nuclear YAP shown on a heatmap) and MFI of active nuclear YAP in EpSC following LTA topical application at different doses (as outlined in E). Median with Standard Error (SE) shown, simple linear regression fitted. **(G)** Representative IF images (dotted line epidermal-dermal boundary, nuclear active YAP shown on a heatmap) and MFI of nuclear YAP in EpSC following LTA topical application at different doses with addition of daily iCOX1 inhibitor. **(H)** Summary Schematic: dose-responsive microbial tuning of COX-1 in LCs results in production of PGE_2_ which modulates EpSC output and promotes barrier integrity. For (A-G): each dot corresponds to data from one mouse, n ≥ 3 mice per condition, median shown, One-Way ANOVA with Šidák multiple-comparison correction.

To shed more light on the underlining mechanism of COX-1 upregulation in LCs we performed two simple experiments. We first tested whether COX-1 induction required sustained microbial exposure. Hence, we compared *S. epi* acute application (1-day) to a longer *S. epi* regime (3-day) and observed that short-term exposure was not sufficient to increase COX-1 expression (Fig. 5C). We then tested whether heat-killed *S. epi* (*S. epi*-HK) could upregulate COX-1 expression, and found that its effect was comparable to live *S. epi,* indicating that a heat-stable component of *S. epi* may be sufficient to induce COX-1 expression in LCs (Fig. 5D).

Focusing on the heat-stable component lipoteichoic acid (LTA) of the gram-positive bacterial cell surface as a potential source, we found that topical LTA on its own caused upregulation of COX-1 protein and *Ptgs1* in LCs (fig. S7A). Although not studied previously in the context of LCs, LTA was particularly interesting in that in inflamed tissues, it can act as a pathogen-associated molecular pattern (PAMP) to elicit inflammatory effects, including prostaglandin release (*54*). These findings therefore suggested that a conserved microbial sensing pathways traditionally linked to inflammation can be repurposed in steady-state skin to tune homeostatic prostaglandin signaling.

Lastly, we sought to dissect the signaling dynamics driving COX-1 induction. We considered two hypotheses: 1) that COX-1 upregulation occurs in a threshold-dependent, switch-like manner, and 2) that commensal cues elicit a graded, dose-dependent response. To distinguish between these possibilities, we topically applied daily doses of LTA for 6 days at varying concentrations, and observed a dose-dependent, linear upregulation of COX-1 in the epidermis (Fig. 5E). Concomitantly, we observed a linear downregulation of nuclear YAP within the EpSC compartment (Fig. 5F).

This reciprocal relation between COX-1 induction and YAP suppression exposed the existence of a rheostat-like mechanism, in which incremental changes in microbial input are translated by LCs into proportional adjustments in the EpSC compartment. Moreover, inhibition of the circuit following iCOX-1 and LTA co-administration resulted in an unstable shift—with increased nuclear YAP activation at higher doses (Fig. 5G) and marked changes in barrier gene expression (fig. S7B-C). Altogether these data reveal a microbiota-immune-epithelial stem cell circuit, underscoring a tight interaction between microbial tuning of COX-1 activity in LCs, followed by their production and release of short-range acting PGE_2_ to govern the balance between EpSC self-renewal and epidermal barrier formation.

## DISCUSSION

As the body’s barrier to the outside world, the epidermis is constantly exposed to environmental stresses—including dehydration, ultraviolet radiation, temperature fluctuations, friction, pathogens, chemicals and mechanical injury. Despite experiencing such diverse perturbations, the skin epithelium has a remarkable capacity to restore and maintain equilibrium, through mechanisms that have remained poorly understood. One way for biological systems to counteract these challenges is to implement control circuits, especially negative feedback, which can dampen noise and buffer against perturbations (*6*).

Here we identified one such circuit, showing that specialized resident macrophages of the epidermis—Langerhans Cells— act as a local rheostat to tightly regulate EpSC renewal. Our data show that LCs bridge communication between EpSCs, safeguarded beneath the barrier they sustain, and the microbiota that colonize the skin’s outermost layers. Through the commensal-tuned, sustained local production of PGE_2_, LCs maintain EpSC quiescence and in so doing, regulate differentiating cell production that leads to refueling the skin barrier. Our data point to the view that, shaped by the microbiota, LCs regulate homeostatic PGE_2_ production in the skin to match EpSC output needs with local microenvironmental conditions.

A biological rheostat such that as the one we unearthed in the current study, is advantageous over a binary on/off control in that it can fine-tune cellular activity in a graded, proportional manner. Such a system can dynamically adjust tissue output to environmental fluctuations while filtering noise and avoiding excessive responses. We speculate that by avoiding excessive EpSC activation, this regulatory strategy may help to preserve stemness while still meeting the need to constantly rejuvenate the skin’s barrier. By positioning this regulatory function within Langerhans cells, their long cellular processes are well suited to integrate signals from the commensal microbiota and pathogenic threats, enabling adaptive communication with epidermal stem cells to tune epidermal flux in response to barrier status.

LCs have recently been shown to influence barrier fitness by supporting the production of antibodies necessary to regulate the local microbial biomass (*55*). Compared to these slower, adaptive immune functions, LC-mediated COX-1 regulation provides a more rapid, temporally distinct mechanism to control EpSCs. Our study extends LC’s functional repertoire from professional antigen-presenting cells (*10, 12*) to central gatekeepers of epidermal barrier integrity. Although there are many possible ways and contexts by which LCs interact with EpSCs, our genetic manipulations of the LC–EpSC niche connection underscore the functional importance of LCs in balancing EpSC self-renewal, quiescence and barrier function formation during skin homeostasis.

The circuit we uncovered places prostaglandin signaling at the interface between microbial influence, immune regulation and stem cell control within the epidermis. Indeed, beyond their classical inflammatory roles, it is appreciated that prostaglandins serve as potent modulators of stem cell fate, integrating environmental and metabolic cues to fine-tune renewal and differentiation, as described for the hematopoietic and intestinal stem cell niches (*56–61*). However, it is only by interrogating the role of prostaglandins in different tissues that it has become apparent that despite signaling through the same EP4 receptor, the downstream pathways engaged in different stem cell compartments diverge markedly, underscoring the context-dependent nature of EP4 signaling, a feature also shared by other prostanoid receptors (*23*).

Indeed our study here stands out in that in addition to uncovering a hitherto unappreciated role for LC:EpSC communication within the epidermis, we identify a central role for the skin microbiota as an upstream regulator of this circuitry. Considering the importance of COX-1 mediated prostaglandins in the gastric mucosa (*23*), it is tempting to speculate that local prostaglandin production might be a shared mechanism to maintain barrier integrity at vulnerable tissue junctions. To this end, recent work has identified a feedback loop in which microbial metabolites activate Tuft-2 cells in the intestine to release prostaglandins, thereby promoting mucus secretion by Goblet cells (*62*). Although the cellular components differ from our epidermal model, the observation that commensal signals induce local prostaglandin production to strengthen barrier function points to the possibility of a potentially conserved, prostaglandin-mediated regulatory mechanism to maintain barrier integrity.

In closing, our study reveals an unexpected convergence of microbiota, resident intra-epithelial macrophages and EpSC control in maintaining and rejuvenating skin integrity. By linking microbial cues to LC-mediated prostaglandin regulation of EpSCs, the epidermis achieves a form of adaptive homeostasis—one that can respond to environmental changes while preserving barrier balance. This work broadens the concept of immune surveillance to include rheostatic control of tissue stem cell output, suggesting that analogous circuits may operate across other barrier surfaces to preserve organ resilience.

## Acknowledgments

We thank L. Polak, L. Hidalgo, I. Crawley, T. Omelchenko, R. Fleming and J. Racelis for technical support. We thank W. Song, N. Guzzi, and Y. Yang, and all member of the Fuchs lab for discussions and manuscript comments. In addition, we thank H. Liu and M. Roulis for critical insights into PG biology. We thank the following investigators for animals and reagents provided (highlighted in the methods): J. Segre (NIH), M. Colonna (Washington University St Louis), D. Mucida (Rockefeller University). FACS was conducted by the Rockefeller University Flow Cytometry Core (S. Svetlana, Director). RNA-seq and scRNA-seq was conducted by the Rockefeller University Genomics Core (C. Zhao, Director); bioinformatics support was provided by Rockefeller University Bioinformatics Core (T. Carroll, Director); and all mouse work was performed in Rockefeller University’s Comparative Bioscience Center and under AAALAC-accreditation according to NIH guidelines for animal care and use.

## Author contribution

Conceptualization: A.G. and E.F. Methodology: A.G., A.R.B., and E.F. Investigation: A.G., R.S., E.G.R., M.S., M.T.T., N.J.A., K.AU.G., S.M.S. Formal analysis: A.G., R.S., E.G.R., M.D.A., L.S.U. Visualization: A.G., R.S., E.G.R., M.S., M.D.A., L.S.U. Validation: A.G., R.S., E.G.R., M.S., M.D.A., L.S.U. Data curation: A.G., R.S., E.G.R., M.S., M.D.A., L.S.U. Funding acquisition: A.G. and E.F. Project administration: A.G. and E.F. Supervision: E.F. Writing – original draft: A.G. and E.F. Writing – review & editing: all authors.

## Data and materials availability

To review bulk RNAseq data, GEO accession GSE312854; Enter token yvutwsaclhijbqd To review scRNAseq data, GEO accession GSE299457

## Supplementary Materials

### The PDF file includes

Materials and Methods

Table S1

Table S2

Supplementary References

Figures and Figure Legends S1 to S7

## MATERIALS AND METHODS

### Animals

C57BL6/J, Rosa26^LSL-Cas9-EGFP^ (JAX:028555), ROSA26^eGFP-DTA^ (JAX:032087) were purchased from The Jackson Laboratory at 6-weeks of age then housed in specific-pathogen-free conditions at the Association for Assessment and Accreditation of Laboratory Animal Care International (AAALAC)-accredited Comparative Bioscience Center at the Rockefeller University. *Il34*^LacZ/LacZ^ were kindly provided by M. Colonna (*1*). huLangerin^CRE/+^ (strain 01X66) came from NCI frozen embryo depository (*2*). K14^CreER/+^ were generated and maintained in house (*3*). Germ-free animals were kindly provided by D. Mucida at Rockefeller University germ-free facility. Unless specified, mice used in the study were in second telogen 7-week-old females or males (no sex difference was observed). Mice were allocated to groups at random and littermate controls were used (although no specific randomization was performed), and appropriate sample sizes were chosen from previous experiments and/or by assessing the known literature; no blinding was performed. All animals were housed in accordance with the procedures delineated in the Guide for the Care and Use of Laboratory Animals. Mice were maintained under a 12-hour light/dark cycle and were provided with food and water *ad libitum*.

### Animal treatments

The following pharmacological inhibitors were topically applied for 24hrs, all suspended in EtOH at 1mg/ml concentration: Indomethacin (S1723), highly selective COX-1 inhibitor SC-560 (S6686), EP4-receptor antagonist ONO-AE3-208 (E2986), 15-PGDH inhibitor SW033291 (S7900) all from Selleckchem, and dmPGE_2_ (Cayman chemicals, Cat: 14750). Lipoteichoic acid (Sigma, L2515) was topically applied daily in EtOH at the described concentrations for 6 days and samples were harvested on day 7. For commensal colonization, *Staphylococcus epidermidis* (strain NIHLM087, obtained from J. Segre) was applied as previously described (*4, 5*). In short, *S. epi* was grown in TBS broth for 18hr, re-suspended in PBS and each mouse was associated with a bacterial suspension (∼10^9^ colony-forming unit [CFU]/mL) across the entire back skin using a sterile cotton swab topically applied to murine back skin every other day using cotton swab for a total of 3 times. For heat killed application, *S. epi* was heat-killed at 95C for 10min (plated to confirm absence of live bacteria). For AA application, 0.1mg of AA-alkyne or AA (Cayman chemical, Cat: 10538 and 90010 respectively) was topically applied to murine back skin for 1hr prior to tissue harvest. Krt14^CreER/+^; Rosa26^LSL-Cas9-EGFP^ animals were injected intraperitoneally with tamoxifen (75 mg/kg tamoxifen in corn oil, both from Sigma-Aldrich) on each of 5 consecutive days and tissue was harvested 5 days post-injection.

### Barrier Assays

Inside-out and outside-in barrier assays were performed using Lucifer Yellow CH, Potassium Salt (ThermoFisher, 100mM stock). For inside-out: euthanized mice were injected subcutaneously with 1mM Lucifer Yellow CH, and incubated at 37C for 30min. For outside-in: euthanized mice had 1mM Lucifer Yellow CH topical application by placing solution in a 24-well plate and dipping the mouse back skin evenly inside, and subsequently incubated at 37C for 1hr (insuring that the skin was constantly covered by solution). Skin samples were collected, washed in PBS and fixation and staining was performed as outlined.

### Organoids

EpSC organoid cultures were made as previously described (*6*). In short, EpSC lines were established in 2D cultures, plated on mitomycin-inactivated 3T3/J2 feeder fibroblast cells, and maintained in E intermediate (300 μM) supplemented with penicillin/streptomycin and 10 μM Y-27632 (Selleckchem) (*7, 8*). Cultures were incubated at 37°C and 7.5% CO2. Media was changed every 2 to 3 days. When near-confluent, EpSCs were digested with 0.25% trypsin-EDTA in PBS (Gibco) for 10 min at 37°C and resuspended with culture media for passaging. For 3D culture, EpSCs were resuspended at a density of 50 to 300 cells/μl in 40 to 75 μl 60% Matrigel (Corning, Cat: 356231) droplets under low-attachment conditions (Nuclon Sphera, Thermo Scientific). After polymerization, cell droplets were initially grown in E300+Y medium supplemented with 50 ng/ml EGF (Peprotech) for 2 to 3 days to maximize colony forming efficiency and outgrowth. Serum-replete culture media was washed with sterile PBS and changed to Dulbecco’s modified eagle media/Ham’s F12 media (DMEM/F12) (3:1) containing 1x B-27 supplement (1:50; Gibco) (84), 300 μM calcium chloride, and 40 mg/l cholesterol-cyclodextrin (Sigma). Pharmacological treatments were performed at days 3-7 by adding either 0.25 µM iEP4 (ONO-AE3-208; Selleckchem Cat: E2986), 0.1 μM dmPGE_2_ (Cayman chemicals, Cat: 14750), 1 μM Verteporfin (Sigma, SML0534), and 10 μM TRULI (The Rockefeller University Lats Inhibitor, kindly provided by Hudspeth lab) (*9, 10*) as outlined in figure legends and organoids were subsequently harvested for IF imaging.

### PKA Colorimetric Activity Assay

PKA Colorimetric Activity Assay Kit (ThermoFisher, Cat: EIAPKA) was performed following kit instructions. In short, keratinocytes were plated in 6-well plate to confluency in regular E intermediate low calcium (50 μM) media (*7*). Subsequently they were overnight serum starved in E intermediate low calcium media without cholera toxin (which constitutively activated PKA) overnight and the next day, dmPGE_2_ (Cayman chemicals, Cat: 14750) or Veh Ctrl was added at stated concentrations for 10min. Samples were then harvested and kept on ice as instructed. Plate was read at 450nm with wavelength correction using Biotech Cytotec 5 plate reader

### Lentivirus production

After sequence verification, plasmids were packaged into a lentivirus by calcium phosphate transfection into 293TN cells with helper plasmids pMD2.G and pPAX2 (Addgene plasmids 12259 and 12260). Viral supernatant was collected 46 hr after transfection and filtered through a 0.45-μm filter. Viral supernatant was concentrated using Pierce™ Protein Concentrator PES, 100K MWCO (Thermo) and centrifuged at 3300 x g.

### *In utero* lentiviral delivery

High-titer lentivirus was prepared and E9.5 embryos of indicated genotypes were infected with lentivirus delivered by ultrasound-guidance microinjection into the amniotic sac as previously described (*11*). At E9.5 the surface ectoderm exists as a single layer of unspecified skin progenitors, which can be efficiently, selectively and stably transduced by the viral DNA, without transduction of dermal cell types. In brief, Cas9 guides selected by VBC.core (Table S2) were cloned into the plasmid sequence (Addgene; plasmid no. 57823 (*12*)) containing a ubiquitous phosphoglycerate kinase promoter (Pgk)–driven Tag-RFP to identify transduced cells. After viral production, 1 μl of either lentivirus was injected in utero into E9.5 Krt14^CreER/+^; Rosa26^LSL-Cas9-EGFP^ embryos. In this system, delivered cargo is stably integrated into the skin epithelial chromosomal DNA and propagated into adulthood in the epidermis.

### Histology - haematoxylin and eosin (H&E)

Skin was fixed with 4% Bouin’s fluid (Electron Microscopy Sciences) in PBS, washed three times for 15 min with PBS and stored in 70% ethanol. Samples were paraffin embedded, sectioned (0.8mm) and stained with haematoxylin and eosin (H&E) by Histowiz Inc.

### Immunofluorescence and microscopy

Murine back skin was fixed for 1hr at room temperature (RT) in 4% paraformaldehyde (Electron Microscopy Sciences) in PBS. Tissue was washed 3X in PBS and switched to 30% sucrose overnight at 4°C. Sections were embedded sagittaly in OCT, and cryosectioned at 20um. Tissue sections were then permeabilized and blocked with 0.1% Triton X-100 and 1X BD Perm/Wash solution in PBS for 45 minutes at RT, and stained with primary antibodies (overnight at 4°C), washed and stained with secondary fluorescence conjugated antibodies (2hr at RT). List of antibodies and clones used can be found in Table S1. AA-alkyne was detected using Click-&-Go Plus 388 Imaging Kit (Vector laboratories). Nuclei were stained with DAPI. Images were acquired on an inverted Dragonfly 202 spinning disk confocal system (Andor Technology Inc.) using the 40x or 63x oil immersion objective and a Zyla camera. Four laser lines (405, 488, 561 and 625 nm) were used for near simultaneous excitation of DAPI, Alexa-448, RRX and Alexa-647 fluorophores. Tiled images with a 1-μm stack step were acquired using the Andor Fusion software (v 2.3), and subsequently stitched with Fusion software.

### Image analysis

Image analysis was performed using Imaris v9.5. Epidermal thickness was determined by KRT14 (basal layer of epidermis) and KRT10 (differentiated payer of epidermis) immunofluorescence. EpSC proliferation was calculated as percent of EdU positive DAPI stained nuclei of total DAPI nuclei within the basal layer (calculated via spot creation in Imaris software) of the epidermis as marked by positive ITGA6 staining. Mean Fluorescence Intensity (MFI) was calculate via surface rending of channel of interest and average MFI calculated. For all, at least 3 images/condition/sample were quantified and averaged for final measurement obtained. For outside-in barrier assay, ImageJ Plot Profile was used to obtain fluorescence spectral profile of Luciferase Yellow-CH channel in the epidermis.

### Fluorescence-activated cell sorting and analysis

Interfollicular epidermal stem cells were isolated as previously described (*13*). To obtain single-cell suspensions for FACS, back skin was dissected and subjected to chemical and mechanical digestion. Telogen skin (was dissected and scraped with a dull scalpel to remove excess fat prior to incubation in 0.25% trypsin/EDTA (GIBCO) for 30 min at 37°C. After trypsinization, cells were released by scraping skin with a dull scalpel on the epidermal side. The resulting cell suspensions were quenched with FACS buffer (5% FBS in PBS), filtered with 40 μm strainers, spun down at 500 x g at 4°C, and washed before incubating with primary antibodies for 30 min on ice. Cells were stained with antibodies (live dead aqua, CD49f, Sca-1, CD200) in 100ul FACS buffer (PBS contained 5% FBS and 1% HEPES). EpSCs were FACS isolated on positive expression of CD49f, Sca-1 and CD200^-^. Lineage negative dump included CD45, CD117, CD140a and CD31. For cell isolation for RNA-seq, cells were sorted on a SONY MA900 sorter with a 100 μm nozzle into RLT buffer, and RNA was extracted using RNeasy Mini Kit as per instruction manual (all from QIAGEN).

### HSC LV transduction and adoptive transfer

Hematopoietic Stem Cells were LV transduced as previously described (*14, 15*). In short, LSKs were obtained from Langerin^Cre/+^; ROSA26^CAS9-eGFP^ animals via enrichment using negative selection Direct Lineage Cell Depletion Kit (Miltenyi, Cat: 130-110-470) using LS columns (Miltenyi) as per kit instructions. Following isolation, LSKs were cultured overnight in SFEM media (Stem cell technologies) with the addition of the following growth factors (final concentrations listed) Flt3 (50 ng/ml), SCF (50 ng/ml), IL7 (10 ng/ml), IL6 (50 ng/ml) and IL3 (10 ng/ml), (all from Peprotrech) with the addition of LV. After 3 days in cultures, cells were sorted on SONY MA900 based on TagRFP expression. 100,000 LSKs were adoptively transferred in irradiated Langerin^Cre/+^; ROSA26^eGFP-DTA^ animals with support bone marrow from Langerin^Cre/+^; ROSA26^eGFP-DTA^ animals and re-constitution of the LC compartment was allowed to ensure for two months post transfer.

### PGE_2_ Elisa

Mouse back skin was flash-frozen with liquid nitrogen, and the epidermis was collected by scrapping with a scalpel blade in a cooled chamber (cryostat chamber was used). Epidermal tissue was placed in a pre-cooled Eppendorf tube with 500ul of PBS together with 1mM EDTA, 15μM Indomethacin, and 15μM SW033291 (to avoid degradation or further production; Selleckchem, Cat: S1723 and S7900 respectively). Samples were then pulverized using sonication beads, spun down at 4C and supernatants collected for downstream analysis. PGE_2_ levels were measured using PGE_2_ Assay as kit instruction (R&D Systems, Cat: KGE004B). Plate was read at 450nm with wavelength correction using Biotech Cytotec 5 plate reader. In the case of topical Arachidonic acid (AA) application, we applied 0.1mg AA (Cayman chemical, Cat: 90010) in EtOH to mouse back skin, and harvested tissues after 1hr for IF fixation or PGE_2_ Elisa.

### RT-qPCR

Total RNA was purified using the RNeasy Mini Kit (QIAGEN) as per instruction manual. DNase I treatment was done on the column for 15 min at room temperature. Quality and concentration of RNA samples were determined using an Agilent 2100 Bioanalyzer. Approximately equivalent amounts of RNA were reverse-transcribed by Superscript VILO reverse transcriptase (Thermo Fisher Scientific). cDNAs were normalized to equal amounts using primers for Ppib. cDNAs were mixed with gene-specific primers and SYBR green PCR Master Mix (Sigma), and qRT-PCR was performed on an QuantStudio6 Flex Real-Time PCR System (ThermoFisher).

### RNA-seq

10,000-20,000 EpSCs per condition were FACS-sorted directly into room temperature buffer RLT (QIAGEN) and immediately flash frozen. Two biological replicates were used per condition, with 2 mice pooled per biological replicate. Total RNA was isolated with RNeasy Mini Kit (QIAGEN) as per manufacturer’s instructions. Low-input RNA libraries were then prepared using Illumina TruSeq low-input mRNA library kit and sequenced on NextSeq 2000 using 50-bp paired-end reads (100 million reads). Transcript quantification was performed using Salmon (v1.10.3) (*16*) in mapping-based mode. The reference index was constructed using the GENCODE vM37 comprehensive transcript annotations, with the entire GRCm39 primary genome assembly included as decoys to perform selective alignment. Quantification was run with the --gcBias flag enabled to correct for fragment-level GC content biases. Transcript-level abundance estimates were aggregated to gene-level counts using the tximport R package (*17*). Differential gene expression analysis was performed using DESeq2 (v1.42.1) (*18*) with default parameters. To improve effect size estimation, log2 fold changes were shrunk using the apeglm method (*19*) for the contrast comparing the *gPtger4*-EpSCs versus the *gCtrl* group. Functional enrichment analyses were performed using the clusterProfiler R package (*20*). For Gene Set Enrichment Analysis (GSEA), genes were ranked based on shrunken log2 fold changes, and the analysis was performed on a specific a priori defined pathway using 10,000 permutations (nPermSimple = 10000) to ensure stability. Nominal p-values were reported without multiple testing correction given the targeted nature of the single-pathway interrogation. A priori gene set for b-Catenin was obtained from GSEA-MSIGDB (GOMF_BETA_CATENIN_BINDING); YAP pathway was curated from known literature (*21*); and CREB signature was obtained from CREB Target Gene Database (*22*). For Over-Representation Analysis (ORA), pathway gene sets were retrieved from the Reactome subset (C2:CP:REACTOME) of the Molecular Signatures Database (MSigDB) for Mus musculus using the msigdbr package. The analysis was focused on significantly upregulated genes (adjusted P value < 0.05 and log2 fold change > 0). To minimize selection bias, the background gene universe was strictly defined as all genes with a detection signal of baseMean > 100 in the DESeq2 analysis. Pathway enrichment was assessed using a hypergeometric test with Benjamini-Hochberg adjustment for multiple comparisons (adjusted P value < 0.05).

### Single-cell RNA-sequencing analyses

Our single cell dataset was compiled by integrating published data (BioProject ID PRJNA625279 at the NCBI SRA database)(*23*) with in-house generated 10X Genomics sequencing data on sorted epidermal stem cells (EpSCs) from untreated C57B6 adult mice (manuscript in press, manuscript in press, GSE299457, only the D30_CTRL_1 and D30_CTRL_2 data were considered for analysis here).

Single cell analysis was performed in R 4.4.1, broadly following the Seurat vignette (Seurat version 5.2.1) and employing the scCustomize package 3.0.1 for visualizing final plots. In more detail, separate SeuratObjects (min.cells = 3, min.features = 200) were created from the filtered feature/barcode/matrix files from PRJNA625279 and the annotated .h5ad input file from in-house generated data. Objects were merged but kept separate before integration to perform preprocessing and quality control. Cells were filtered based on the following metrics: minimal number of 200 genes, maximal number of 8000 genes, a mitochondrial percentage of less than 15% to exclude dead cells. Data were normalized using LogNormalize, variable features were identified using the “vst” function (nfeatures = 3000). Cells were subjected to CellCycleScoring and doublets were identified and subsequently removed using r-scrublet, for which we loaded the python package scrublet(*24*) (https://github.com/AllonKleinLab/scrublet.git) within a conda environment in R. After quality metrics were completed, the two datasets were integrated based on reference anchors as previously identified using the SelectIntegrationFeatures and FindIntegrationAnchors functions in Seurat. Data scaling was performed on the integrated data, regressing out potentially confounding variables, which included mitochondrial percentage, doublet and cell cycle scores as well as RNA count per cell. Following the general Seurat vignette, we performed principal components analysis (PCA) on our integrated data, determining dimensionality by calculating percent of variation per PC, cumulative percent for each PC and limiting the number of PCs to those, whose change in percent of variation between subsequent PCs exceeded 0.1%. With these PC dimensionality values at hand, clustering and UMAP visualization were performed as per Seurat guidelines and UMAP clustering resolution was reviewed and refined using clustree 0.5.1. Clusters were named based on the expression of top features per cluster. Epithelial cells (EpSCs and HFSCs) from PRJNA625279 were included when running data integration to provide an overlap in cell types with our in-house dataset and thus enable proper data integration. For final analyses, however, the EpSC and HFSC cluster from PRJNA625279 were excluded, the integrated data were re-scaled followed by PCA scoring and UMAP re-clustering as described above. The original cluster IDs were maintained, though. Final Dimplots, Featureplots, Violinplots and Dotplots were created using the scCustomize package 3.0.1.

For differential gene expression (DEG) analysis, done on the “RNA” assay (normalized data) of our integrated dataset, we grouped all immune cell clusters except for Langerhans cells and compared them to Langerhans cells using the FindMarkers function in Seurat. Due to off-the-scale plotting of highly significant datapoints on a conventional VolcanoPlot, we plotted DEG data on an MA plot – plotting the average expression level (x-axis) versus the log-fold change (y-axis) of DEGs - using the ggplot2 3.5.1 package.

For receptor:ligand analyses we used CellChat (*25*) (version 2.1.2), created a CellChat Object from our integrated SeuratObject, and followed the general CellChat vignette as described (https://htmlpreview.github.io/?https://github.com/sqjin/CellChat/blob/master/tutorial/CellChat-vignette.html). We customized the CellChat receptor:ligand database by manually curating receptor:ligand pair entries associated with Prostaglandin signaling, which were missing in the original database. Given that prostaglandins constitute secreted lipids that would not be represented in transcriptional data, we used their synthesizing enzymes and annotated receptors as proxies. In summary, we included the following pairs: PTGS1:PTGER1-4, PTGS:PTGER1-4, PTGDS:PTGDR, PTGDS:PTGDR2, PTGES:PTGER1, PTGES2-3:PTGER1, PTGES:PTGER2, PTGES2-3:PTGER2, PTGES:PTGER3, PTGES2-3:PTGER3, PTGES:PTGER4, PTGES2-3:PTGER4. The normalized RNA data of our integrated dataset were used as input for all receptor:ligand analyses via CellChat.

### Statistics

Statistical test were performed in Prism software v10. The statistical tests used, and annotations for significance are provided for each figure/figure legend. Group sizes for EpdSC proliferation were determined by power analysis on the basis of preliminary experimental results whenever possible. For all data shown, data is pooled from at least 2-independent experiments. For all genomics experiments, the replicate numbers, read depth and quality control assessments followed the ENCODE Consortium’s guidelines and best practices (https://www.encodeproject.org/data-standards/). Manuscript was checked by Proofig AI.

## SUPPLEMENTARY MATERIALS

Materials and Methods

Table S1

Table S2

Supplementary Figures and Supplementary Figure Legends, S1 to S7

Supplementary References

---

**Table S1:** List of Antibodies for IF.

| Name | Species | Ab clone | Company |
| --- | --- | --- | --- |
| KRT14 | Chicken | Polyclonal 9060 | Biologend |
| KRT10 | Rabbit | Polyclonal 19054 | Biologend |
| Langerin | Rat | 929F3.01 | Novus |
| ITGA6 | Rat | GoH3 | Biologend |
| BrdU | Mouse | MoBU-1 | Invitrogen |
| Survivin | Rabbit | Polyclonal | Cell Signaling |
| COX-1 | Rabbit | Polyclonal | Cell Signaling |
| RFP | Goat | Polyclonal | MyBioSource |
| YAP | Rabbit | EPR19812 | abcam |
| FILAGRIN | Rabbit | Lab generated | Fuchs Lab |
| LORICRIN | Rabbit | Lab generated | Fuchs Lab |
| TGM1 | Rabbit | Polyclonal | Novus |
| KRT6 | Rabbit | Polyclonal 19057 | Biologend |
| F4/80 | Rat | BM8 | Biologend |

**Table S2:**
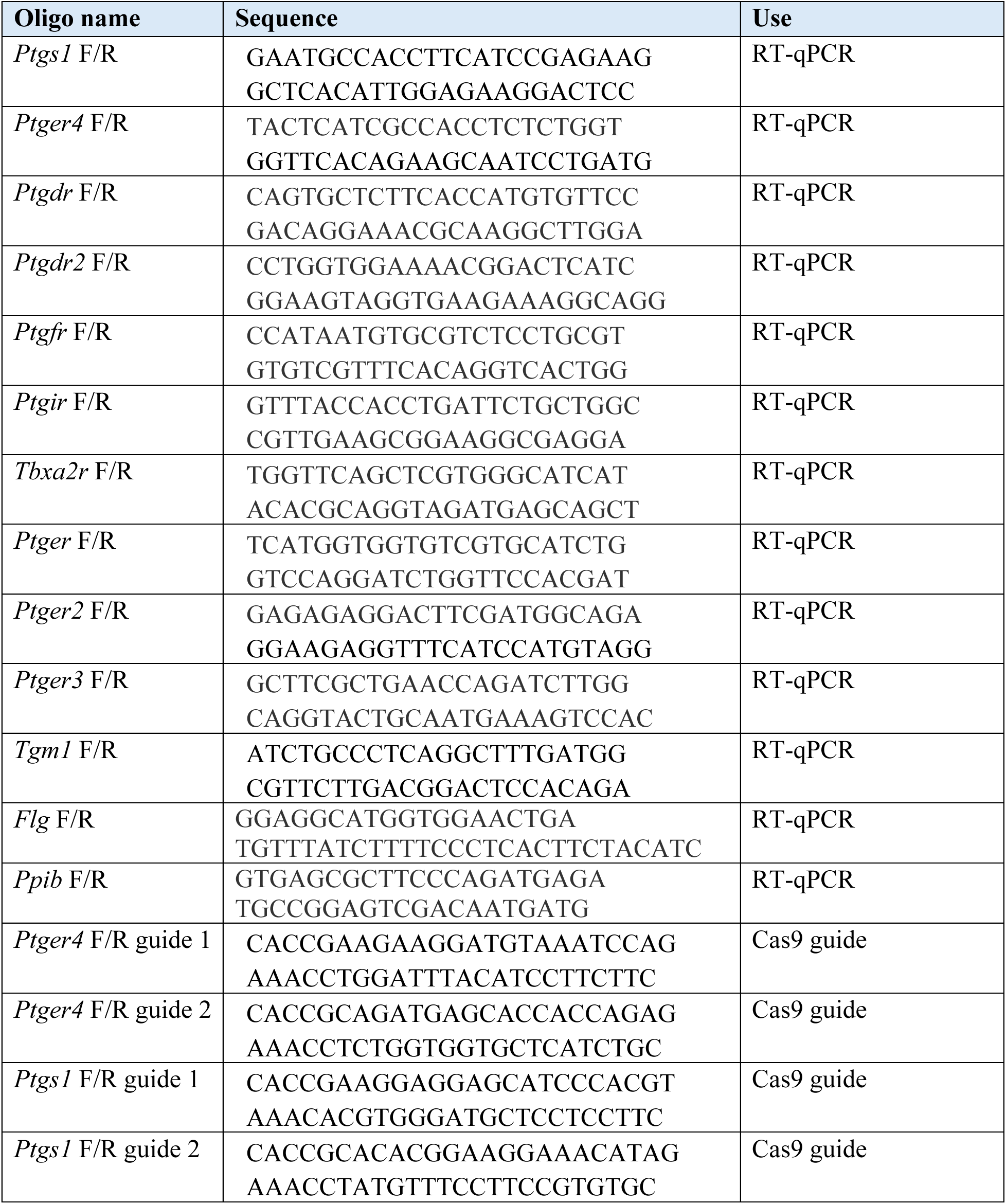
Oligo primer sequences.

**Supplementary Figure 1:**
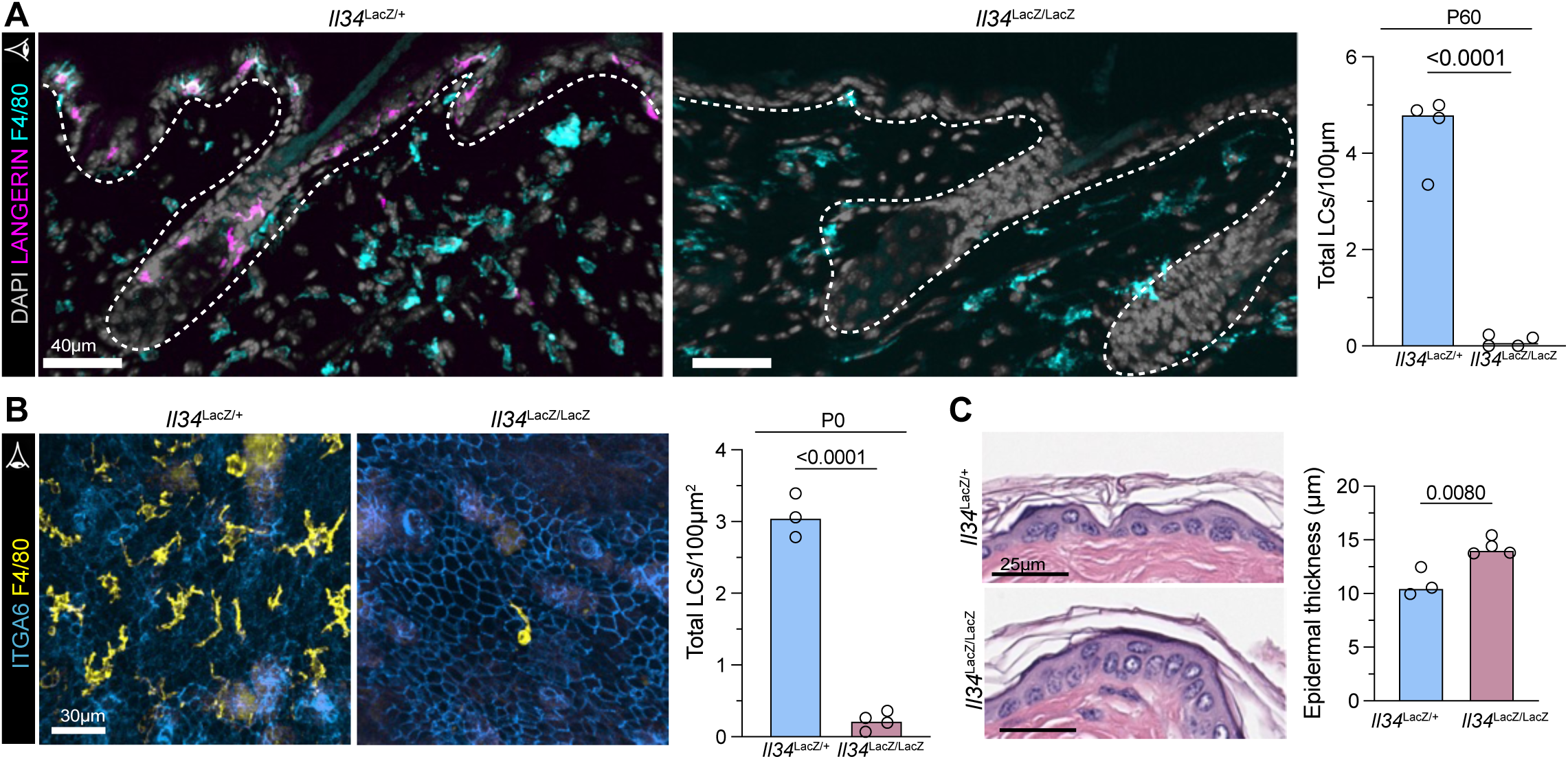
Characterization of *Il34*^LacZ/LacZ^ animals. **(A)** Representative IF images and quantifications of LC numbers in control and LC-deficient animals (*Il34*^LacZ/+^ and *Il34*^LacZ/LacZ^ respectively) adult (P60) animals. Dashed line denotes epidermal-dermal boundary; no other F4/80^+^ cell can compensate in the epidermis for the absence of LCs in *Il34*^LacZ/LacZ^ animals. **(B)** Representative IF images (top-down view) and quantifications of LC numbers in *Il34*^LacZ/+^ and *Il34*^LacZ/LacZ^ animals at P0. Of note, Langerin is not expressed by LCs prior to P3, F4/80 is used as an alternative LC marker. **(C)** Representative H&E sagittal sections for *Il34*^LacZ/+^ and *Il34*^LacZ/LacZ^ animals and quantification of epidermal thickness. For (A-C): Each dot corresponds to data from one mouse with median shown, schematic eyes denote sagittal vs planar orientations of sections, n ≥ 3 mice per condition, two-tailed unpaired Student’s t tests.

**Supplementary Figure 2:**
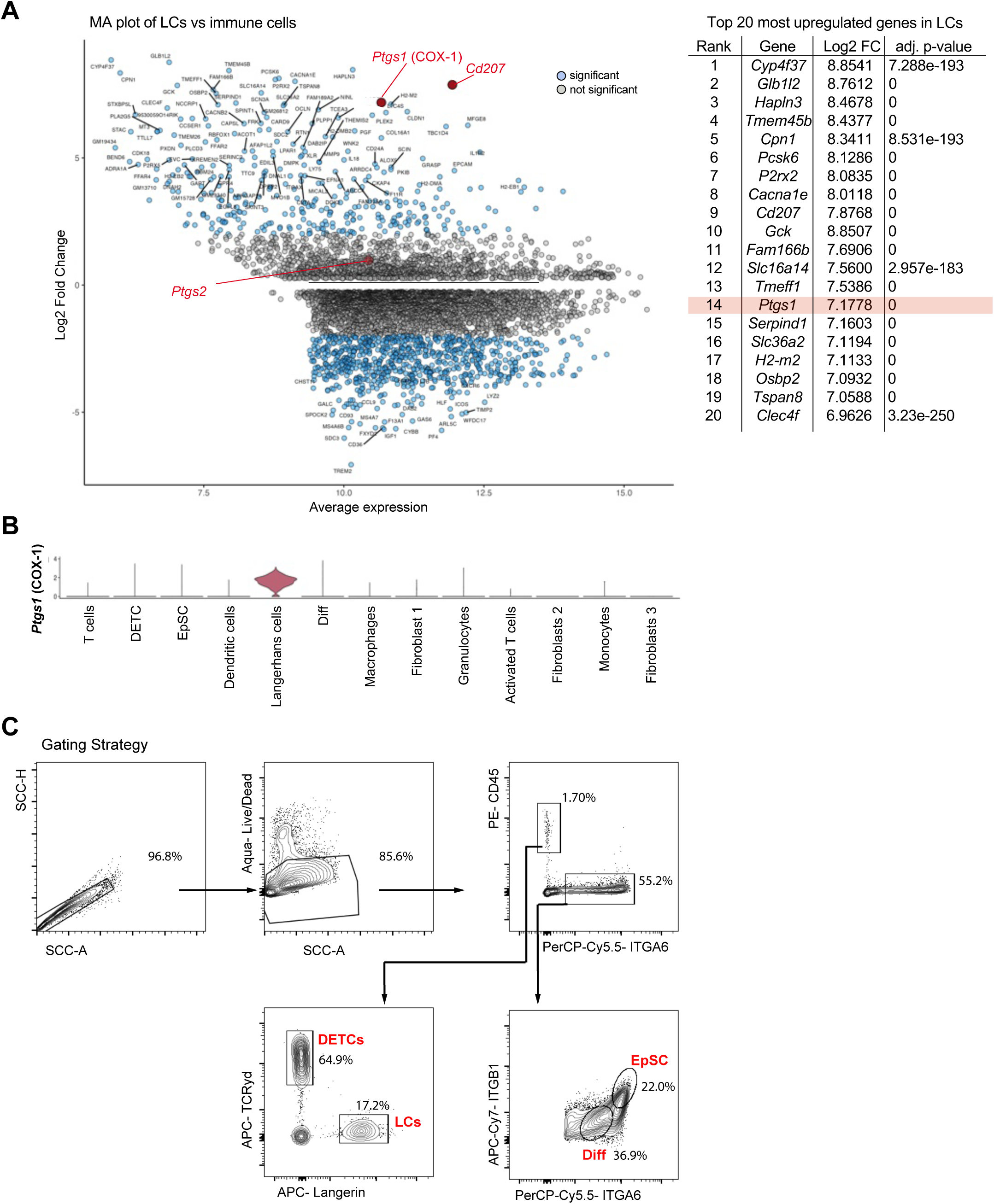
COX-1 is highly enriched in LCs. **(A)** Left: MA plot comparing LCs to all other skin resident immune cells highlighting *Ptgs1* (COX-1) expression in LCs as one of the most abundant and enriched genes. Right: list of top 20 most upregulated genes in LCs, their fold change expression when compared to all other skin-resident immune cells and adjusted p-value. **(B)** Violin plot of *Ptgs1* expression levels in all cell types from single cell RNAseq UMAP in F2. **(C)** Flow cytometry plots depict the gating scheme and relative percentages of EpSCs, Diff, DETCs and LCs populations.

**Supplementary Figure 3:**
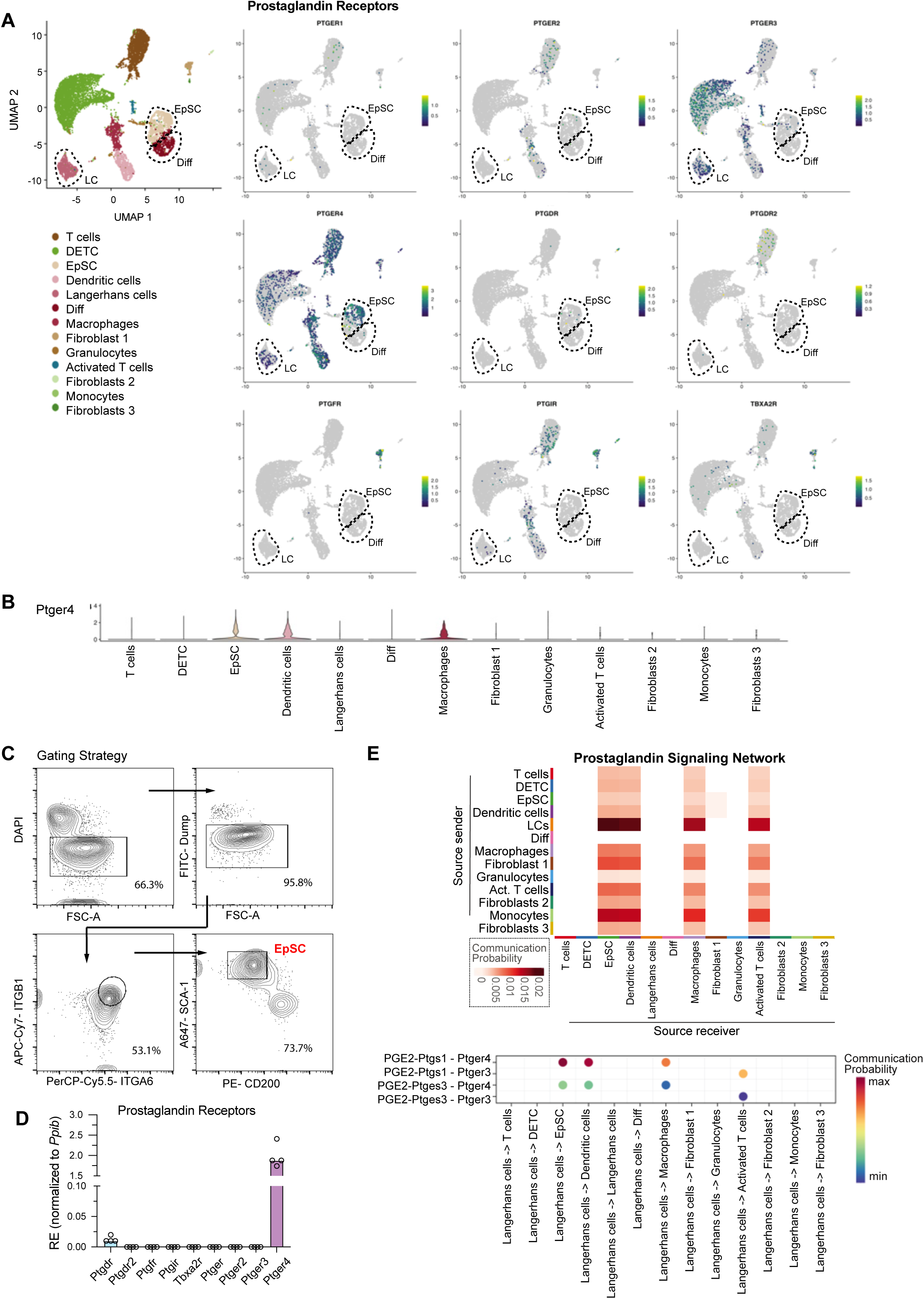
EP4 is the major PG receptor expressed by EpSCs. **(A)** UMAP plots showing the expression of all PGs receptors within the skin present within the dataset. For receptors not listed, no transcripts were detected. **(B)** Violin plot of *Ptger4* expression levels in all cell types from single cell RNAseq UMAP in F2. **(C)** Flow cytometry plots depict sorting strategy for EpSC. **(D)** Sorted EpSCs Relative Expression (RE) of prostaglandin receptor expression (normalized to housekeeping *Ppib*) as quantified by qPCR. **(E)** Top: Communication probability of the Prostaglandin Signaling Network between all cell types with source senders and source receivers as determined by Cell-Chat. Bottom: Cell-Chat analysis highlighting the individual gene pairs enriched in the Prostaglandin Signaling Network.

**Supplementary Figure 4:**
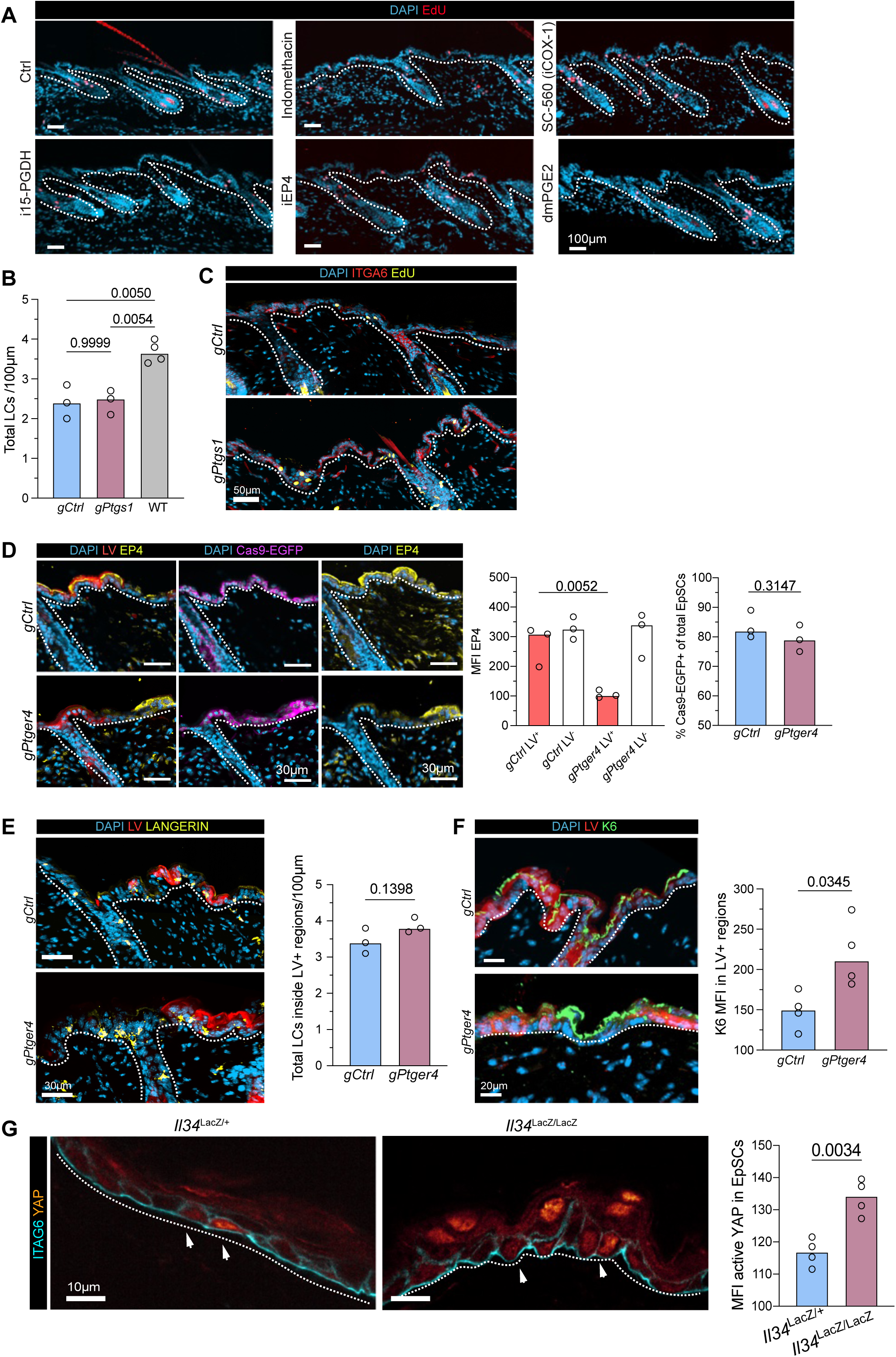
Modulation of PGs in the epidermis results in altered EpSC behavior. **(A)** Representative IF images showing EpSC proliferation (S-phase EdU labeling) following 24h pharmacological PG manipulation (quantification shown in Fig. 3A-B). **(B)** Quantification of total LC numbers in *gCtrl* and *gPtgs1* animals when compared to WT Ctrls; One-Way ANOVA with Šidák multiple-comparison correction. **(C)** Representative IF image showing EpSC proliferation and hyper-thickening in *gCtrl* and *gPtgs1* animals (quantifications shown in Fig. 3F). **(D)** Representative IF image and Mean Fluorescence Intensity (MFI) quantifications of EP4 knockdown (One-Way ANOVA with Šidák multiple-comparison correction), and frequency of Cas9-EGFP expression (two-tailed unpaired Student’s t tests) in *gCtrl* and *gPtger4* animals following TAM administration. **(E)** Representative IF image and quantification of total number of LCs in *gCtrl* and *gPtger4* animals; two-tailed unpaired Student’s t tests. **(F)** Representative IF image and quantification of K6 MFI in LV^+^ regions; two-tailed unpaired Student’s t tests. **(G)** Representative IF image and quantifications of active nuclear YAP (channel shown as a heatmap) in *Il34*^LacZ/+^ and *Il34*^LacZ/LacZ^; two-tailed unpaired Student’s t tests. For (A-G), each dot corresponds to data from one mouse, dashed line denotes epidermal-dermal boundary, n ≥ 3 mice per condition, median shown.

**Supplementary Figure 5.**
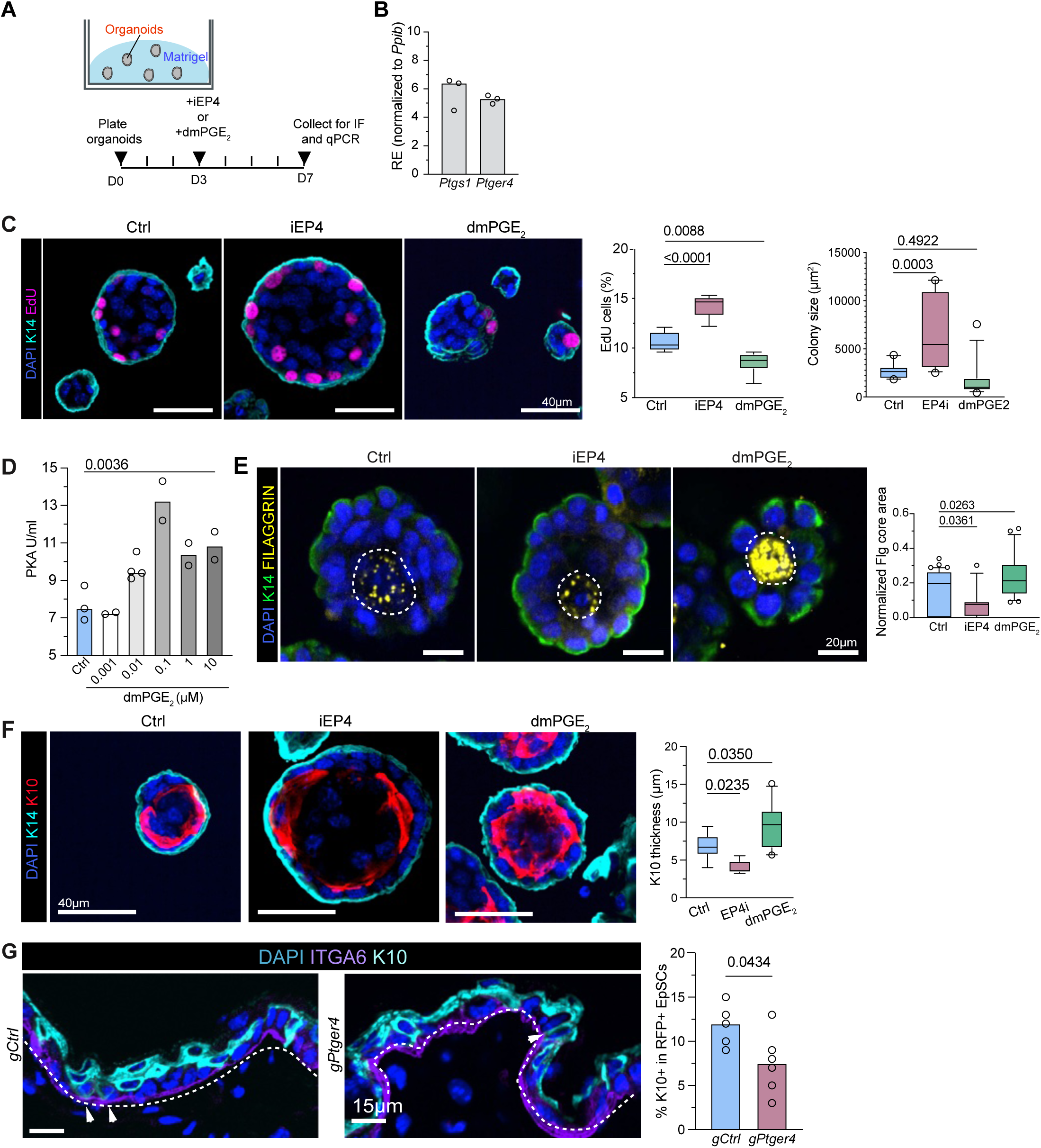
Modulation of PG signaling *in vitro* recapitulates altered EpSC function *in vivo*. **(A)** Schematic of EpSC organoid culture system. **(B)** RE of *Ptgs1* and *Ptger4* in organoids harvested at D3 (normalized to housekeeping *Ppib*). **(C)** Representative IF image of organoids treated with vehicle control, iEP4 or dmPGE_2_ and quantification of colony size, and EpSC proliferation (S-phase EdU labeling). Box-and-Whiskers plot (10-90 percentile); One-Way ANOVA with Šidák multiple-comparison correction. **(D)** Levels of PKA activation in cultured EpSC following dmPGE_2_ administration. Each dot represents a biological replicate (pooled from two technical replicates), One-Way ANOVA with Šidák multiple-comparison correction. **(E)** Representative IF image and quantification FILAGGRIN core area (normalized to total organoid area) in Ctrl, iEP4, and dmPGE_2_ treated organoids. Each dot represents an organoid, One-Way ANOVA with Šidák multiple-comparison correction. **(F)** Representative IF image and quantifications of K10 thickness in Ctrl, iEP4, and dmPGE_2_ treated organoids. Each dot represents an organoid, Box-and-Whiskers plot (10-90 percentile). One-Way ANOVA with Šidák multiple-comparison correction. **(G)** Representative IF images and quantifications of the percentage of committed K10^+^ K14^+^ double-positive out of the total K14^+^ pool of basal layer progenitors in *gCtrl* and *gPtger4* LV^+^ regions, dashed line denotes epidermal-dermal boundary; two-tailed unpaired Student’s t tests, each dot represents one mouse, n ≥ 5 mice per condition, median shown.

**Supplementary Figure 6.**
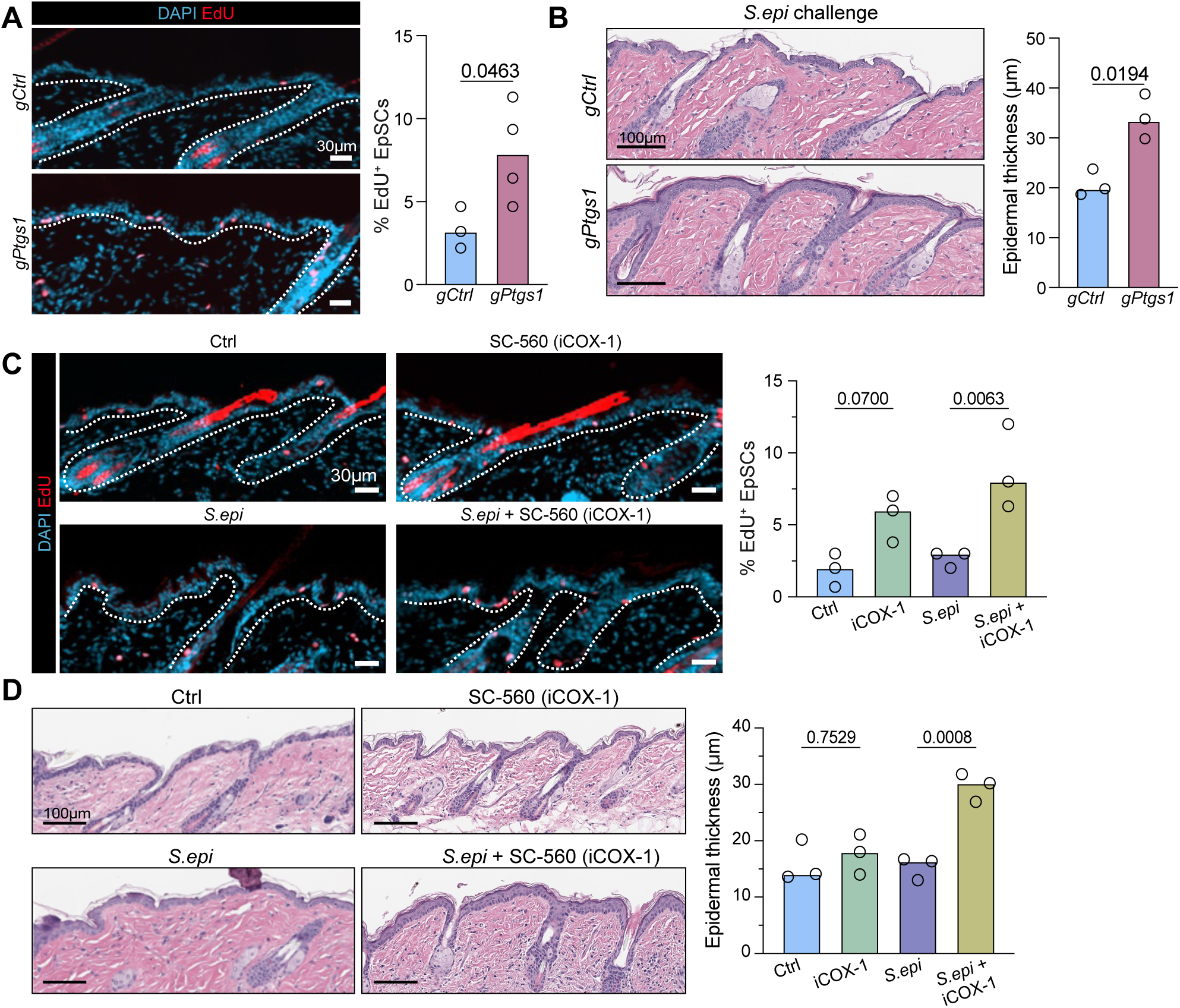
Loss of epidermal barrier function in EP4 deficient EpSCs. Representative IF image and quantifications of EpSC proliferation (S-phase EdU labeling) in *gCtrl* and *gPtgs1* animals following *S. epi* administration; two-tailed unpaired Student’s t tests. (**A**) Representative H&E staining and quantification of epidermal thickness in *gCtrl* and *gPtgs1* animals following *S. epi* administration; two-tailed unpaired Student’s t tests. **(C)** Representative IF image and quantification of EpSC proliferation (S-phase EdU labeling) in daily pharmacologically treated animals concomitant with *S. epi* administration; One-Way ANOVA with Šidák multiple-comparison correction. **(D)** Representative H&E staining and quantification of epidermal thickness in daily pharmacologically treated animals concomitant with *S. epi* administration; One-Way ANOVA with Šidák multiple-comparison correction. For all, each dot corresponds to data from one mouse, dashed line denotes dermal-epidermal boundary, n ≥ 3 mice per condition, median shown.

**Supplementary Figure 7.**
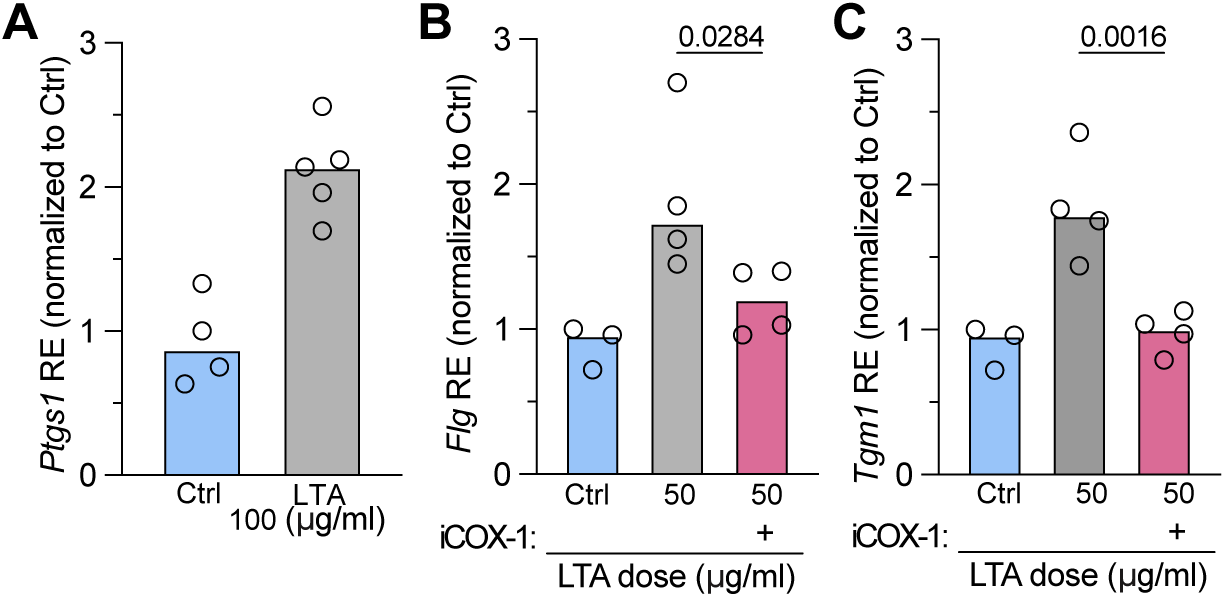
LTA is sufficient to promote COX-1 upregulation and up-regulate barrier gene expression. Epidermal Relative Expression (RE) of *Ptgs1* following LTA (100 μg/ml) daily topical application for 6 days (normalized to untreated Ctrl) as quantified from qPCR; two-tailed unpaired Student’s t tests. **(B-C)** Epidermal RE of *Flg* and *Tgm1* respectively following LTA (50 μg/ml) daily topical application +/- with iCOX-1 inhibitor daily topical application (normalized to untreated Ctrl); One-Way ANOVA with Šidák multiple-comparison correction. For all: each dot corresponds to data from one mouse, n ≥ 3 mice per condition, median shown.

